# The genome of an apodid holothuroid (*Chiridota heheva*) provides insights into its adaptation to deep-sea reducing environment

**DOI:** 10.1101/2021.09.24.461635

**Authors:** Long Zhang, Jian He, Peipei Tan, Zhen Gong, Shiyu Qian, Yuanyuan Miao, Han-Yu Zhang, Qi Chen, Qiqi Zhong, Guanzhu Han, Jianguo He, Muhua Wang

## Abstract

Cold seeps and hydrothermal vents are deep-sea reducing environments that are characterized by a lack of oxygen, photosynthesis-derived nutrients and a high concentration of reducing chemicals. Apodida is an order of deep-sea echinoderms lacking tube feet and complex respiratory trees, which are commonly found in holothurians. *Chiridota heheva* Pawson & Vance, 2004 (Apodida: Chiridotidae) is one of the few echinoderms that resides in deep-sea reducing environments. Unlike most cold seep and hydrothermal vent-dwelling animals, *C. heheva* does not survive by maintaining an epi- or endosymbiotic relationship with chemosynthetic microorganisms. The species acquires nutrients by extracting organic components from sediment detritus and suspended material. Here, we report a high-quality genome of *C. heheva* as a genomic reference for echinoderm adaptation to reducing environments. *Chiridota heheva* likely colonized its current habitats in the early Miocene. The expansion of the aerolysin-like protein family in *C. heheva* compared with other echinoderms might be involved in the disintegration of microbes during digestion, which in turn facilitates the species’ adaptation to cold seep environments. Moreover, several hypoxia-related genes were subject to positive selection in the genome of *C. heheva*, which contributes to their adaptation to hypoxic environments.

## 1. Introduction

Echinodermata is a phylum of marine animals comprising 5 extant classes, including Asteroidea (starfish), Ophiuroidea (brittle star), Echinoidea (sea urchin), Crinoidea (feather star), and Holothuroidea (sea cucumber) (Pawson, 2007). Adult echinoderms are characterized by having a body showing pentameral symmetry, a water vascular system with external tube feet (podia), and an endoskeleton consisting of calcareous ossicles (Pechenik, 2015). Echinoderms exhibit a high divergence in morphology, from the star-like architecture in Asteroidea to the worm-like architecture in Holothuroidea (Mooi & David, 2008; Smith et al., 2013).

Compared with other echinoderms, holothurians have a unique body architecture and evolutionary history. The worm-like body of the holothurian preserves the pentameral symmetry structurally along the oral-aboral axis (Li et al., 2020). In addition, holothurians have a soft and stretchable body, in which the ossicles are greatly reduced in size (Pechenik, 2015). Variations in body architecture also exist in Holothuroidea. The order Apodida is a group of holothurians that are found in both shallow-water and deep-sea environments (Pawson & Vance, 2004). Phylogenetic analyses showed that Apodida is sister to other orders of Holothuroidea (Lacey et al., 2005; Miller et al., 2017). Apodid holothurians lack tube feet and complex respiratory trees, making them morphologically distinct from other holothurians (Pechenik, 2015). In contrast to other classes of Echinodermata, which experienced a severe evolutionary bottleneck during the Permian-Triassic mass extinction interval, Holothuroidea did not experience family-level extinction through the interval. The deposit-feeding lifestyle of holothurians conferred a selective advantage during the primary productivity collapse of the Permian-Triassic mass extinction (Twitchett & Oji, 2005). As the genome of only one shallow-water holothurian (*Apostichopus japonicus*) has been assembled and analyzed (Li et al., 2018; Zhang et al., 2017), it is critical to study the genomes of more holothurians to dissect their evolutionary history and developmental processes.

Cold seeps and hydrothermal vents are deep-sea reducing environments that are characterized by high hydrostatic pressure, low temperature, lack of oxygen and photosynthesis-derived nutrients, and high concentrations of reducing chemicals (Levin, 2005). However, these harsh environments support a variety of macroinvertebrates, including tubeworms, mussels, clams, and gastropods (Vanreusel et al., 2009). Most of these macrobenthos depend on the epi- or endosymbiotic relationships with chemoautotrophic microorganisms for nutrition (Van Dover et al., 2002). Recent genomic analyses have revealed the genetic basis of adaptation in several seep- and vent-dwelling macrobenthos hosting symbiotic bacteria (Li et al., 2019; Sun et al., 2020; Sun et al., 2017; Y. Sun et al., 2021). However, nonsymbiotic animals residing in deep-sea reducing environments are understudied with only one reported genome (Liu et al., 2020).

Hydrocarbon fluid seepage from cold seeps is completely devoid of O_2_ and comprises high levels of sulfides. After reacting with sulfides contained in the fluid, any free O_2_ is removed from the deep-sea water. There are unique mechanisms of hypoxic adaptation in seep-dwelling animals, as their O_2_ consumption rates are similar to those of related shallow-water species (Hourdez & Lallier, 2007). However, the genomic basis of hypoxic adaptation in seep-dwelling animals is still lacking.

Echinoderms are a rare component of deep-sea chemosynthetic ecosystems (Tunnicliffe, 1992). *Chiridota heheva* Pawson & Vance, 2004 (Apodida: Chiridotidae) is one of the few echinoderms that occupies all three types of chemosynthetic ecosystems (hydrothermal vent, cold seep, and whale fall) (Thomas et al., 2020). This suggests that the species is well adapted to deep-sea reducing environments. Unlike most seep- and vent-dwelling species, *C. heheva* does not host chemosynthetic bacteria (Pawson & Vance, 2004). *Chiridota heheva* derives nutrients from a variety of sources, extracting organic components from sediment detritus, suspended material, and wood fragments when available (Carney, 2010; Pawson & Vance, 2004). A peltate-digitate tentacle structure allows *C. heheva* to exploit various food sources by switching between deposit and suspension feeding (Thomas et al., 2020). The cosmopolitan distribution and special lifestyle of *C. heheva* make it an ideal model to study adaptation to deep-sea reducing environments in nonsymbiotic animals.

Here, we assembled and annotated a high-quality genome of *C. heheva* collected from the Haima cold seep in the South China Sea. The evolutionary history of *C. heheva* was investigated by inferring the phylogenetic relationship among echinoderms and the demographic history of *C. heheva* and a shallow-water holothurian (*Apostichopus japonicus*). Additionally, comparative genomic analyses were performed to dissect the genomic basis of adaptation to deep-sea reducing environments in *C. heheva*.

## 2 Methods and Materials

### 2.1 Sample collection and genome sequencing

The *C. heheva* sample used in this study was collected using manned submersible *Shenhaiyongshi* from the Haima cold seep in the South China Sea (16° 73.228’ N, 110° 46.143’ E, 1,385 m deep) on August 2, 2019. The *C. heheva* individuals were kept in an enclosed sample chamber placed in the sample basket of the submersible. Once the samples were brought to the upper deck of the mothership, the muscle of the individuals was dissected, cut into small pieces, and immediately stored at -80°C. The samples were then transported to Sun Yat-sen University on dry-ice and stored at -80°C until use.

To construct Nanopore sequencing library, high molecular weight genomic DNA was prepared by the CTAB method. The quality and quantity of the DNA were measured via standard agarose gel electrophoresis and with a Qubit□4.0 Fluorometer (Invitrogen). Sequencing library was constructed and sequenced by Nanopore PromethION platform (Oxford Nanopore Technologies). Additionally, DNA was extracted to construct Illumina sequencing library. The quality and quantity of the DNA were measured via standard agarose gel electrophoresis and with a Qubit□2.0 Fluorometer (Invitrogen). Sequencing library was constructed and sequenced by Illumina Novaseq platform (Illumina).

### 2.2 Genome assembly

#### 2.2.1 Mitochondrial genome assembly

Low quality (reads with ≥10% unidentified nucleotide and/or ≥50% nucleotides having phred score < 5) and sequencing-adaptor-contaminated Illumina reads were filtered with custom C script. The filtered Illumina reads were then trimmed with Fastp (v0.21.0) (Chen et al., 2018) to obtain high-quality Illumina reads, which were used in the following analyses. Mitochondrial genome of *C. heheva* was assembled using the two-step mode of mitoZ (v2.4) (Meng et al., 2019) with the high-quality Illumina reads. And the assembled genome was annotated using mitoZ (v2.4) with parameter “--clade Echinodermata”.

#### 2.2.2 Nuclear genome assembly

The size and heterozygosity of *C. heheva* genome were estimated using high-quality Illumina reads by *k*-mer frequency distribution method. The number of *k*-mers and the peak depth of *k*-mer sizes at 17 was obtained using Jellyfish (v2.3.0) (Marcais & Kingsford, 2011) with the *-C* setting. Genome size was estimated based on the *k*-mer analysis as described previously (Star et al., 2011).

Low quality Nanopore reads were filtered using custom Python script. Two draft genome assemblies were generated using filtered Nanopore reads with Shasta (v0.4.0) (Shafin et al., 2020) and WTDBG2 (v2.5) (Ruan & Li, 2020), respectively. The contigs of the two draft assemblies were subject to error correction using filtered Nanopore reads with Racon (v1.4.16) (Vaser et al., 2017) three times. The corrected contigs were then polished with high-quality Illumina reads with Pilon (v1.23) (Walker et al., 2014) three times. The error-corrected contigs of Shasta assembly and WTDBG2 assembly were assembled into longer sequences using quickmerge (v0.3) (Chakraborty et al., 2016). The merged contigs were subject to error correction using filtered Nanopore reads with Racon three times, and then using high-quality Illumina reads with Pilon three times. As the heterozygosity of *C. heheva* genome is high, haplotypic duplications in the assembled genome were identified and removed using purge_dups (v1.2.3) (Guan et al., 2020). The completeness and quality of the assembly was evaluated using BUSCO (v4.0.5) (Simao et al., 2015) against the conserved Metazoa dataset (obd10), and SQUAT with high-quality Illumina reads (Yang et al., 2019).

### 2.3 Genome annotation

#### 2.3.1 Repetitive element annotation

Repetitive elements in the assembly were identified by *de novo* predictions using RepeatMasker (v4.1.0) (https://www.repeatmasker.org/). A *de novo* repeat library for *C. heheva* was built using RepeatModeler (v2.0.1) (Flynn et al., 2020). To identify repetitive elements, sequences from the *C. heheva* assembly were aligned to the *de novo* repeat library using RepeatMasker (v4.1.0). Additionally, repetitive elements in *C. heheva* genome assembly were identified by homology searches against known repeat databases using RepeatMasker (v4.1.0). A repeat landscape of *C. heheva* genome was obtained using an R script that was modified from https://github.com/ValentinaBoP/TransposableElements.

#### 2.3.2 Protein-coding gene annotation

We applied a combination of homology-based and *de novo* predication methods to build consensus gene models for the *C. heheva* genome assembly. For homology-based gene prediction, protein sequences of *Helobdella robusta, Lytechinus variegatus, Strongylocentrotus purpuratus, Dimorphilus gyrociliatus, Apostichopus japonicus* and *Acanthaster planci* were aligned to the *C. heheva* genome assembly using tblastn. The exon-intron structures then were determined according to the alignment results using GenomeThreader (v1.7.0) (Gremme et al., 2005). In addition, *de novo* gene prediction was performed using Augustus (v3.3.2) (Stanke et al., 2006), with the parameters obtained by training the software with protein sequences of *Drosophila melanogaster* and *Parasteatoda tepidariorum*. Two sets of gene models were integrated into a non-redundant consensus gene set using EvidenceModeler (v1.1.1) (Haas et al., 2008). To identify functions of the predicted proteins, we aligned the *C. heheva* protein models against NCBI NR, trEMBL, and SwissProt database using blastp (E-value threshold: 10^−5^), and against eggNOR database (Huerta-Cepas et al., 2019) using eggNOR-Mapper (Huerta-Cepas et al., 2017). In addition, KEGG annotation of the protein models was performed using GhostKOALA (Kanehisa et al., 2016).

### 2.4 Phylogenomic analysis

Protein sequences of 15 metazoan species (*A. planci, S. purpuratus, Lytechinus variegatus, A. japonicus, Anneissia japonica, Saccoglossus kawalevskii, Branchiostoma floridae, Ciona intestinalis, Danio rerio, Gallus gallus, H. robusta, Mus musculus, Pelodiscus sinensis, Petromyzon marinus*, and *Xenopus laevisproteins*) were downloaded from NCBI. And protein sequences of *Parastichopus parvimensis* were downloaded from Echinobase (Kudtarkar & Cameron, 2017). OrthoMCL (v2.0.9) (Li et al., 2003) was applied to determine and cluster gene families among these 16 metazoan species and *C. heheva*. Gene clusters with >100 gene copies in one or more species were removed. Single-copy othologs in each gene cluster were aligned using MAFFT (v7.310) (Katoh et al., 2002). And the alignments were trimmed using ClipKit (v1.1.3) (Steenwyk et al., 2020) with “gappy” mode. The phylogenetic tree was reconstructed with the trimmed alignments using a maximum-likelihood method implemented in IQ-TREE2 (v2.1.2) (Minh et al., 2020) with *H. robusta* as outgroup. The best-fit substitution model was selected by using ModelFinder algorithm (Kalyaanamoorthy et al., 2017). Branch supports were assessed using the ultrafast bootstrap (UFBoot) approach with 1,000 replicates (Hoang et al., 2018).

To estimate the divergent time among echinoderms, single-copy orthologs were identified among *A. japonica, A. planci, A. japonicus, P. parvimensis, C. heheva, L. variegatus* and *S. purpuratus* after running OrthoMCL pipeline as mentioned above. Single-copy orthologs were aligned using MAFFT (v7.310), trimmed using ClipKit (v1.1.3) with ‘gappy’ mode, and concatenated using PhyloSuite (v1.2.2) (Zhang et al., 2020). Divergent time among 7 echinoderms were estimated using the concatenated alignment with MCMCtree module of the PAML package (v4.9) (Tessmar-Raible & Arendt, 2003). MCMCtree analysis was performed using the maximum-likelihood tree that was reconstructed by IQ-TREE2 as a guide tree and calibrated with the divergent time obtained from TimeTree database (minimum = 193 million years and soft maximum = 350 million years between *L. variegatus* and *S. purpuratus*).

### 2.5 Demographic inference of *C. heheva* and *A. japonicus*

Paired-end Illumina reads of *A. japonicus* (Li et al., 2018) were downloaded from NCBI SRA database. The reads of *A. japonicus* were filtered with custom C script and trimmed with fastp (v0.21.0). The Illumina clean reads of *C. heheva* and *A. japonicus* were aligned to the respective reference genome assembly using BWA (v0.7.17) (Li & Durbin, 2009) with “mem” function. Genetic variants were identified using Samtools (v1.9) (Li et al., 2009). Whole genome consensus sequence was generated with the genetic variants using Samtools (v 1.9). PSMC (v0.6.5) (Li & Durbin, 2011) was used to infer the demographic history of *C. heheva* and *A. japonicus* using the whole genome consensus sequences. The substitution mutations rate and generation time of *C. heheva* and *A. japonicus* was set to 1.0 × 10^−8^ and 3 years according to the previous study of *A. planci* (Hall et al., 2017).

### 2.6 Homeobox gene analysis

Homeobox genes in *C. heheva* genome were identified by following the procedure described previously (Marletaz et al., 2019). Homeodomain sequences, which were retrieved from HomeoDB database (http://homeodb.zoo.ox.ac.uk) (Zhong et al., 2008), were aligned to *C. heheva* genome assembly using tbalstn. Sequences of the candidate homeobox genes were extracted based on the alignment results. The extracted sequences were aligned against NCBI NR and HomeoDB database to classify the homeobox genes.

### 2.7 Gene family evolution

#### 2.7.1 Gene family expansion and contraction analysis

r8s (v1.7) (Sanderson, 2003) was applied to obtain the ultrametric tree of 7 echinoderm species, which is calibrated with the divergent time between *A. planci* and *S. purpuratus* (541 mya). CAFÉ (v5) (De Bie et al., 2006) was applied to determine the significance of gene family expansion and contraction among 7 echinoderm species based on the ultrametric tree and the gene clusters determined by OrthoMCL (v2.0.9).

#### 2.7.2 Evolutionary analysis of *C. heheva* NOD-like receptors (NLRs) and other representative metazoan NLRs

We used HMMER (v3.1) to search against the proteome of *C. heheva* with the HMM profile of NACHT domain (PF05729) retrieved from Pfam 34.0 as the query and an *e* cut-off value of 0.01. Proteins identified by the HMM search were retrieved from the proteome and aligned with 964 representative proteins from eukaryotes and prokaryotes (Urbach & Ausubel, 2017), and other representative metazoan NLRs (Yuen et al., 2014) using hmmalign method implemented in HMMER (v3.1) based on the STAND NTPase domain. The alignment was refined by manual editing. The large-scale phylogenetic analysis was performed using an approximate maximum likelihood method implemented in FastTree (Price et al., 2010). Representative SWACOS and MalT NTPases were used as outgroups (Urbach & Ausubel, 2017). Significant hits clustering with metazoan NLRs were regarded as NLRs, and protein domain organizations were annotated through hmmscan method implemented in HMMER (v3.1).

To explore the evolutionary relationships among *C. heheva* NLRs and other representative metazoan NLRs, we reconstructed the phylogenetic tree of NLRs. The NACHT domains of *C. heheva* NLRs and representative metazoan NLRs were aligned using MAFFT (v7.310), and then refined by manual editing. The representative metazoan NLRs were chosen from literature (Yuen et al., 2014). The phylogenetic tree was reconstructed using a maximum-likelihood method implemented in IQ-TREE2 (v2.1.2). The best-fit substitution model selected by using ModelFinder algorithm. Branch supports were assessed using the UFBoot approach with 1,000 replicates.

### 2.8 Identification and analysis of positively selected genes

Branch-site models implemented in the codeml module of the PAML package is widely used to identified positively selected genes (PSGs). Thus, we identified PSGs in the *C. heheva* genome within the single-copy orthologs among 7 echinoderm species, based on the branch-site models using GWideCodeML (v1.1) (Macias et al., 2020). *C. heheva* was set as the ‘foreground’ phylogeny, and the other species were set as the ‘background’ phylogeny. An alternative branch site model (Model = 2, NSsites = 2, and fix_omega = 0) and a neutral branch site model (Model = 2, NSsites = 2, fix_omega = 1, and omega = 1) were tested. Genes with Bayesian Empirical Bayes (BEB) sites > 90 % and a corrected *P*-value < 0.1 were identified to have been subject to positive selection.

To investigate *LHPP* gene evolution, sequences of *LHPP* from 8 mammals (*Odobenus rosmarus, Orcinus orca, Lipotes vexillifer, Tursiops truncates, Physeter catodon, Balaenoptera acutorostrata, Mus musculus*, and *Homo sapiens*) and 7 echinoderms (*A. japonica, A. planci, A. japonicus, P. parvimensis, C. heheva, L. variegatus* and *S. purpuratus*) were aligned using MAFFT (v7.310). To reconstruct the phylogenetic tree, OrthoMCL (v2.0.9) (Li et al., 2003) was applied to determine and cluster gene families among these 15 species. Gene clusters with >100 gene copies in one or more species were removed. Single-copy othologs in each gene cluster were aligned using MAFFT (v7.310) (Katoh et al., 2002). And the alignments were trimmed using ClipKit (v1.1.3) (Steenwyk et al., 2020) with “gappy” mode. The phylogenetic tree was reconstructed with the trimmed alignments using a maximum-likelihood method implemented in IQ-TREE2 (v2.1.2) (Minh et al., 2020). *H. robusta* was used as outgroup. The best-fit substitution model was selected by using ModelFinder algorithm (Kalyaanamoorthy et al., 2017).

## 3. Results

### 3.1 Characterization and genome assembly of *C. heheva*

The sequenced sample was collected at a depth of 1,385 meters using manned submersible *Shenhaiyongshi* from the Haima cold seep in the South China Sea (16° 73.228’ N, 110° 46.143’ E) (**Figure 1**). We sequenced the sample genome on the Nanopore and Illumina sequencing platforms. In total, 42.43 Gb of Nanopore reads and 49.19 Gb of Illumina reads were obtained (**Table S1 and S2**). Species identity of the sequenced individual was first determined according to its morphological characteristics. In addition, we assembled the mitochondrial genome of the individual using Illumina reads. The sequence identity between the published *C. heheva* mitochondrial genome and our assembled genome was 99.74%, which confirmed the species identity of the sequenced individual (S. Sun et al., 2021). Based on the *k*-mer distribution of Illumina reads, the size of the *C. heheva* genome was estimated to be 1.23 Gb with a high heterozygosity of 2% (**Figure S1 and Table S3**). The *C. heheva* genome was assembled into 4,609 contigs, with a total size of 1.107 Gb and contig N50 of 1.22 Mb (**Table 1**). We determined the completeness of the assembled genome by running benchmarking universal single-copy orthologs (BUSCO) and sequencing quality assessment tool (SQUAT) software. BUSCO analysis with metazoan (obd10) gene set showed that the assembled *C. heheva* genome contained 89.6% complete single-copy orthologs (**Table S4**). Additionally, 91.1% of Illumina reads could be aligned to the assembled genome with high confidence in SQUAT analysis (**Table S5**). These results indicate the high integrity of our assembled genome.

**Table 1.** Genome assembly statistics of *C. heheva* and *A. japonicus*.

|  | *C. heheva* | *A. japonicus*<br>(Li et al., 2018) | *A. japonicus*<br>(Zhang et al., 2017) |
| --- | --- | --- | --- |
| **Estimated genome size (Gb)** | ~1.23 | ~1.0 | ~1.0 |
| **Assembled genome size (bp)** | 1,106,937,276 | 952,279,490 | 804,993,085 |
| **Number of contigs** | 4,609 | 21,303 | 7,058 |
| **Contig N50 (bp)** | 1,221,604 | 45,411 | 190,269 |
| **Scaffold N50 (bp)** | - | 195,518 | 486,650 |

**Figure 1.**
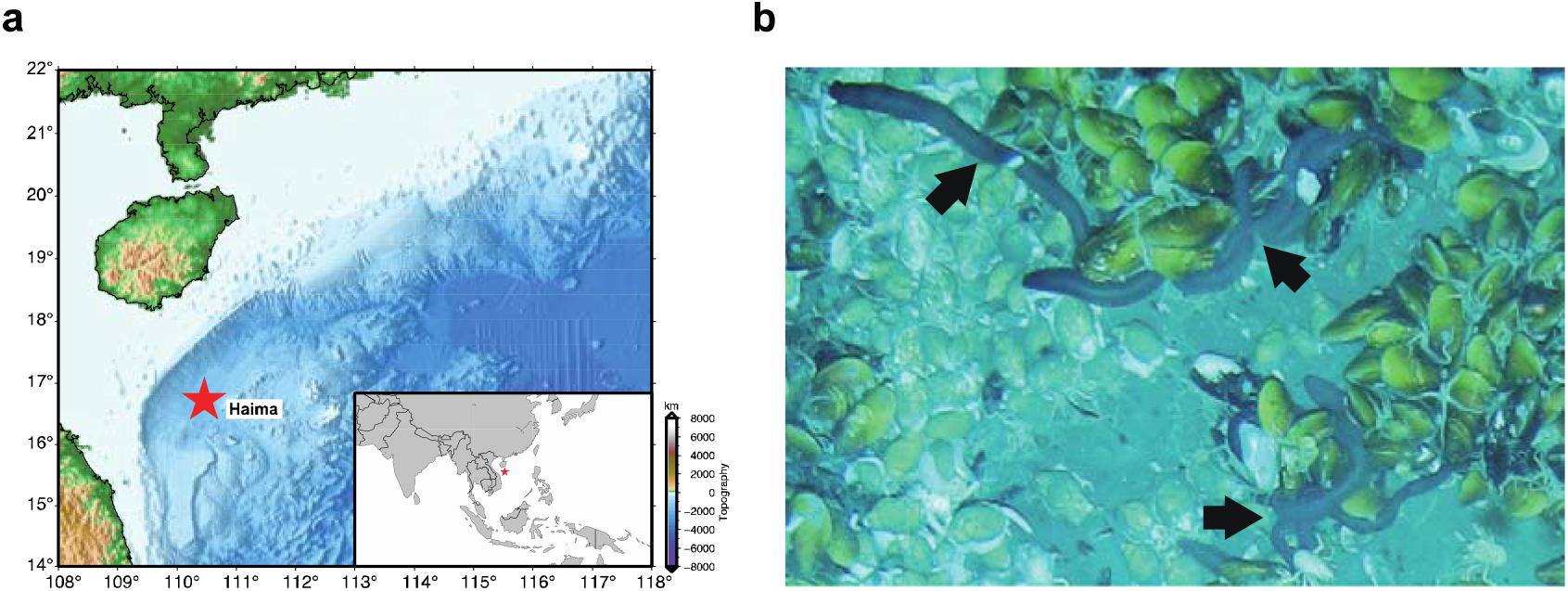
Collection of *C. heheva*. (a) Map showing the sampling site at the Haima cold seep of northern South China Sea (16° 73.228’ N, 110° 46.143’ E). (b) *C. heheva* at the sampling site (depth: 1,385 m), where they cohabit with deep-sea mussels. *C. heheva* individuals are indicated by black arrows.

### 3.2 Genome annotation

Repetitive elements represented 624.38 Mb (56.40%) in the *C. heheva* genome assembly (**Table S6**). Long interspersed nuclear elements (LINEs) were the largest class of annotated transposable elements (TEs), making up 9.72% of the genome. DNA transposons, which were the second largest class of TEs, represented 33.59 Mb (3.03%) of the genome. Additionally, the *C. heheva* genome comprised a large proportion (38.39%) of unclassified interspersed repeats. Comparative genomic analysis among *C. heheva* and other echinoderms revealed that the *C. heheva* genome consisted of the largest number of TEs (**Figure 2a and 2b; Table S7**). Repetitive elements constituted 624.38 Mb of the *C. heheva* genome, and they accounted for 253.98 Mb and 218.2 Mb of the genomes of *A. japonicus* and *P. parvimensis*, respectively. The differences in the repeat content were almost consistent with the size differences between the genomes of *C. heheva* and the other two holothurians. This suggests that repeats contributed to the size differences among the genomes of these three holothurians. Notably, the proportion of LINEs in the *C. heheva* genome was substantially higher than that in the genomes of other echinoderms (**Figure 2b**). Kimura distance-based copy divergence analysis identified a recent expansion of LINEs in the *C. heheva* genome (**Figure 2c**). Protein-coding genes were identified in the genome of *C. heheva* through a combination of *ab initio* and homology-based protein prediction approaches. In total, we derived 36,527 gene models in the *C. heheva* genome. The structure of predicted genes in *C. heheva* is slightly different to that of other previously sequenced echinoderm genomes. With longer exon and intron as well as more exons per gene, genes in *C. heheva* are longer than the ones in *A. japonicus* (**Table S8**).

**Figure 2.**
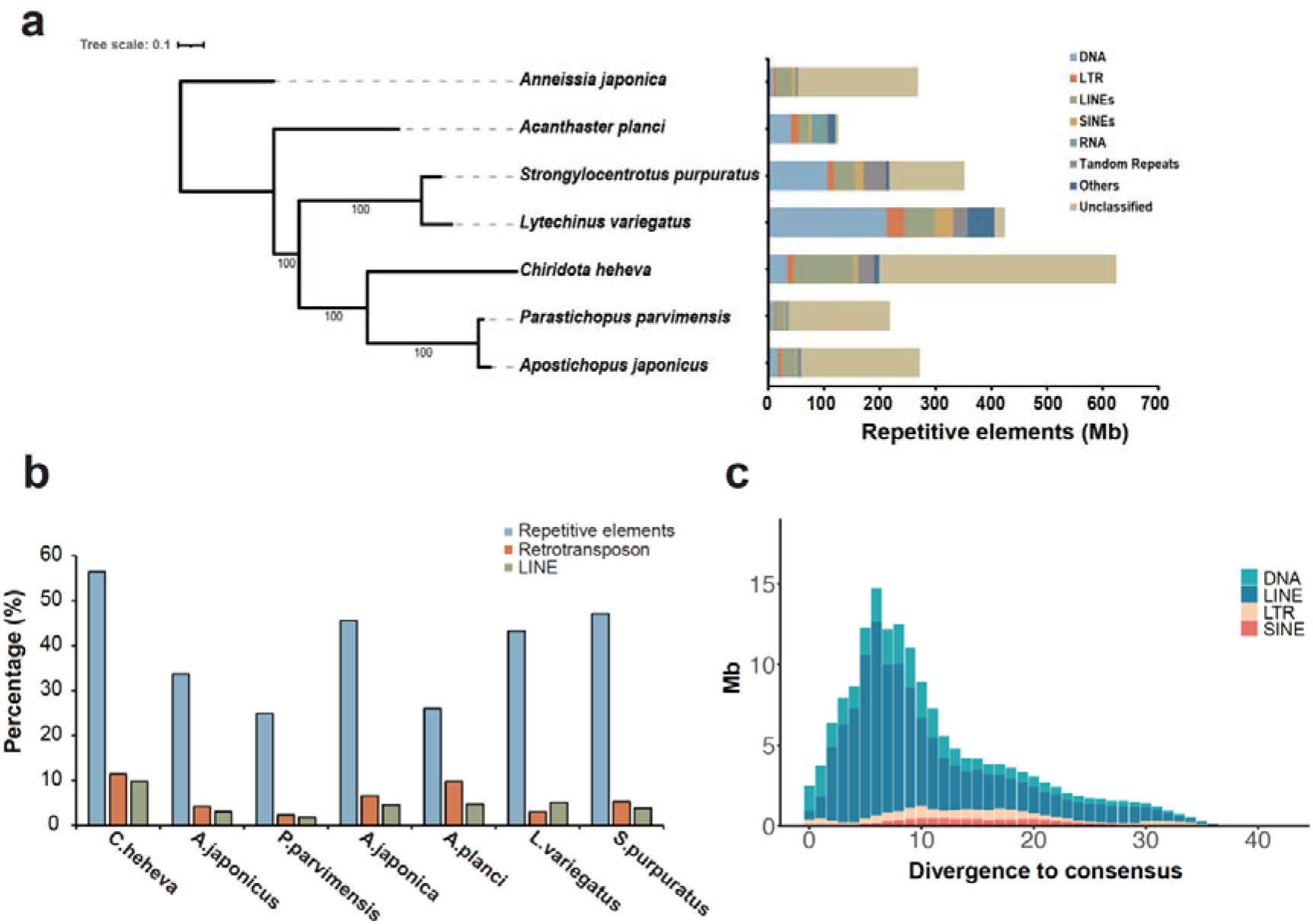
Landscape of transposable elements in echinoderm genomes. (a) Comparison of the occurrence and composition of repetitive elements in the genomes of 7 echinoderms. (b) Comparison of the proportion of repetitive elements, retrotransposon, and long interspersed nuclear elements (LINEs) in the genomes of 7 echinoderms. The proportions of repetitive elements and LINEs are higher in the genome of *C. heheva* than that in other echinoderms. (c) Transposable element accumulation profile in *C. heheva* genome. A recent burst of LINEs was observed in *C. heheva*.

### 3.3 Phylogenomic analysis and demographic inference

To investigate the evolutionary history of *C. heheva*, a maximum-likelihood (ML) phylogenetic tree was reconstructed using single-copy orthologs of *C. heheva* and 16 other deuterostomes (**Figure S2**). Consistent with the results of previous analyses, the tree showed that Echinodermata and Hemichordata were sister groups to Chordata. *Chiridota heheva* appeared sister to two other holothurians, which supports the view that Apodida is the sister taxon to the remaining holothuroids (Miller et al., 2017). In addition, divergence times were determined among 7 echinoderms that had whole genome sequences (**Figure 3a**). The divergence time of *A. japonica* and other echinoderms was estimated to be approximately 569 million years (Ma), which is generally consistent with the fossil records (Smith, 1988a; Zamora et al., 2013). *Chiridota heheva* and two other holothurians were estimated to have diverged approximately 429 Ma. As Apodid is the basal taxon in Holothuroidea, these results indicate that holothurians started to diverge in the Early Ordovician (Benton & Twitchett, 2003).

**Figure 3.**
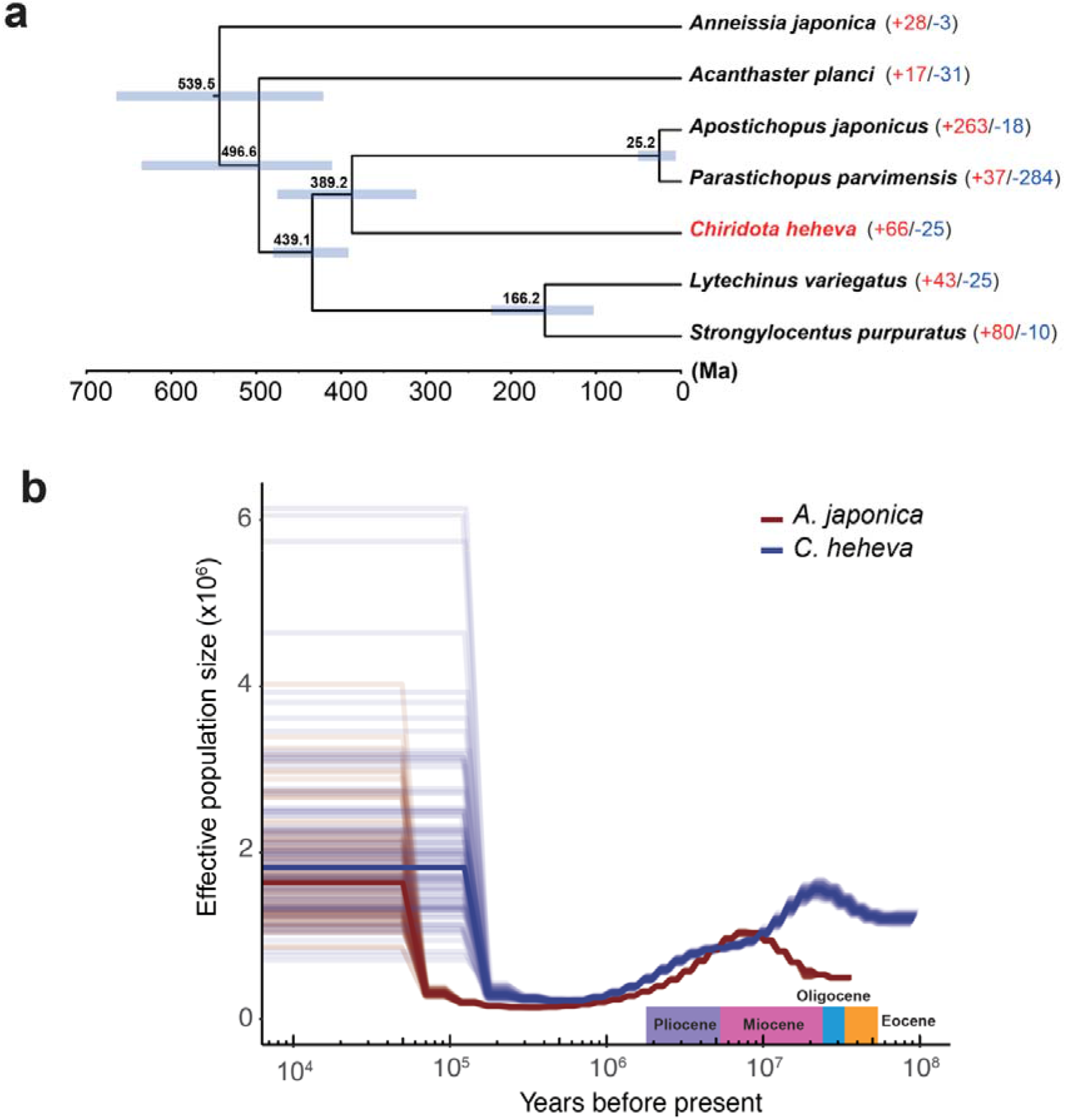
Evolutionary history of *C. heheva*. (a) A species tree of 7 echinoderm species. In total, 988 single-copy orthologs were used to reconstruct the phylogenetic tree. The divergence time between species pairs was listed above each node, and 95% confidence internal of the estimated divergence time was denoted as blue bar. The numbers of protein families that were significantly expanded (red) of contracted (blue) (*P* < 0.05) in each species are denoted beside the species names. (b) Demographic history of *C. heheva* (blue) and *A. japonicus* (red). The changes of ancestral population size of *C. heheva* and *A. japonicus* were inferred using the PSMC method. Time in history was estimated by assuming a generation time of 3 years and a mutation rate of 1.0×10^−8^.

We studied the demographic history of the deep-sea (*C. heheva*) and shallow-water (*A. japonicus*) holothurians by inferring the histories of ancestral population size using the pairwise sequential Markovian coalescent (PSMC) method (**Figure 3b**). *Chiridota heheva* experienced a decline in population size approximately 21 million years ago, which suggests that this species started to colonize the current habitat at the turn of the Miocene. The decline in population size in *A. japonicus* started in the late Miocene (approximately 8 Ma). *Chiridota heheva* also experienced a moderate decline in ancestral population size in the early Pliocene.

### 3.4 *Hox/ParaHox* gene clusters

It has been demonstrated that *Hox* genes play a critical role in embryonic development (Pearson et al., 2005). In addition, previous studies proposed that the presence/absence and expression pattern of *Hox* genes might contribute to morphological patterning and embryonic development in echinoderms (Li et al., 2018; Zhang et al., 2017). Therefore, to determine whether *Hox* genes contribute to morphological divergence in Holothuroidea, we identified *Hox* gene clusters and their evolutionary sister complex, the *ParaHox* gene cluster, in the genomes of *C. heheva* and 6 other echinoderms (**Figure 4**). A *Hox* cluster and a *ParaHox* cluster could be identified in the genomes of all 7 species. The gene composition and arrangement of both *Hox* and *ParaHox* clusters were highly consistent between the genomes of *C. heheva* and *A. japonicus*, suggesting that *Hox/ParaHox* genes do not control the development of tube feet and respiratory trees in Apodida. *Hox4* was missing in Echinodeans and holothurians, and *Hox6* was missing in asteroideans and holothurians. These results support the view that the absence of *Hox* genes might have contributed to the morphological divergence of echinoderms.

**Figure 4.**
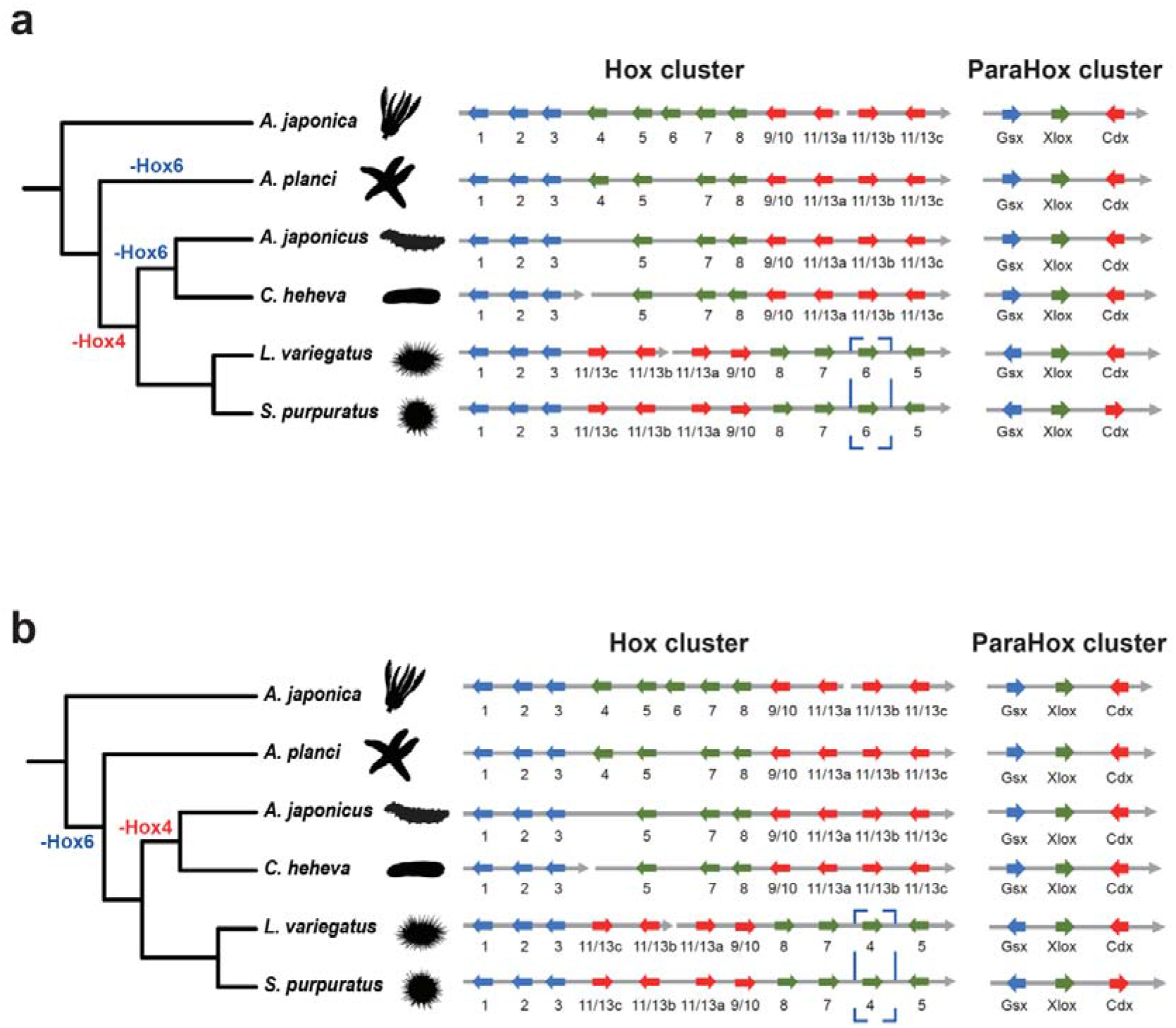
Genomic organization of *Hox* and *ParaHox* gene clusters in echinoderms. *Hox* and *ParaHox* genes are indicated by arrows. The gene composition and orientation of *Hox* and *ParaHox* clusters are consistent between two holothurians (*C. heheva, A. japonicus*). (a) The organization of *Hox* genes in echinoderms if *Hox4* is lost in echinoids (*L. variegatus, S. purpuratus*). (b) The organization of *Hox* genes in echinoderms if *Hox6* is lost in echinoids.

### 3.5 NLR repertoire in *C. heheva*

NACHT and leucine-rich, repeat-containing proteins (NLRs) are important components of pathogen recognition receptors (PRRs) involved in animal innate immune systems, which can perceive pathogen-associated molecular patterns (PAMPs) of viruses and bacteria (Lange et al., 2011). The bona fide NLRs contain a NACHT (NAIP, CIITA, HET-E, and TP1) domain, which belongs to the signal transduction ATPases with numerous domains (STAND) superfamily, and a series of C-terminal leucine-rich repeats (LRRs) (Ausubel, 2005; Leipe et al., 2004). The Pfam hidden Markov model (HMM) search combined with phylogenetic analysis approach identified only 53 NLRs in *C. heheva* (**Table S9**), compared with a largely expanded set of 203 NLRs in purple sea urchin, a member of the phylum Echinodermata (Hibino et al., 2006). *Chiridota heheva* contained 24 NLRs with one or more N-terminal Death/DED domain, 22 NACHT-only NLRs, 6 NLRs with other domains, including the immunoglobulin V-set domain, which was not identified in sea urchin NLRs, and only one NLR with LRRs (**Table S9**). Taken together, these results indicate that the *C. heheva* NLR repertoire shows different abundances and structural complexities than the sea urchin.

We performed phylogenetic analysis of *C. heheva* NLRs and other representative NLRs of metazoans, including humans, *Amphimedon queenslandica, S. purpuratus, Acropora digitifera, Nematostella vectensis, Pinctada fucata, Capitella teleta*, mollusks, and arthropods (Yuen et al., 2014). We found that the majority of *C. heheva* NLRs form a monophyletic lineage with sea urchin NLRs (**Figure 5**), supporting the lineage-specific evolution of NLRs in Echinodermata (Zhang et al., 2010). Given that human IPAF (ice protease-activating factor) and NAIP (neuronal apoptosis inhibitory protein) proteins were reported to have originated before the evolution of vertebrates (Zhang et al., 2010), one *C. heheva* NLR clustering with these two proteins indicates that this NLR may have an ancient independent origin (**Figure 5**).

**Figure 5.**
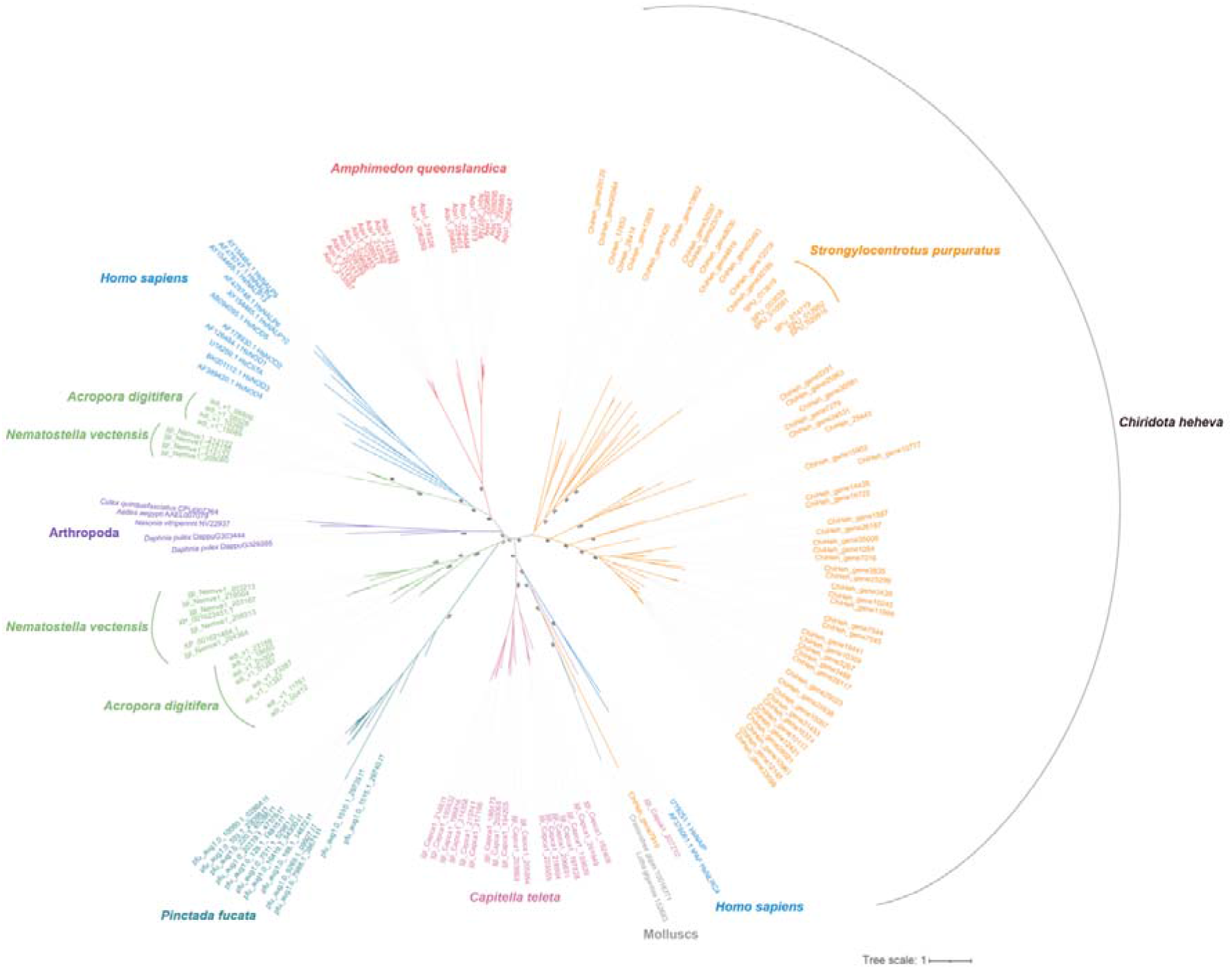
Evolutionary relationships among *C*.*heheva* NLRs and other representative metazoan NLRs. The unrooted phylogenetic tree was reconstructed based on the NACHT domain sequences using a maximum likelihood method. The values near the nodes are ultrafast bootstrap (UFBoot) values. NLRs from different types of species are highlighted by branches of different colors. The species name is shown near the corresponding lineage.

### 3.6 Gene family evolution

We performed gene-family analysis based on the phylogenetic tree of 7 echinoderms (**Figure 3a**). Compared with other echinoderms, 66 gene families were expanded, and 25 gene families were contracted in *C. heheva* (*P* < 0.05) (**Tables S10 and S11**). Several significantly expanded gene families are involved in the processes of cell cycle progression, protein folding, and ribosome assembly. As high hydrostatic pressure causes cell cycle delay and affects protein folding (George et al., 2007; Yancey & Siebenaller, 2015), expansion of these families may have contributed to the adaptation of *C. heheva* to cold seep environments.

Aerolysins, which are pore-forming toxins (PFTs), were first characterized as virulence factors in the pathogenic bacterium *Aeromonas hydrophyla* (Abrami et al., 2000; Dal Peraro & van der Goot, 2016). The homologs of aerolysin in eukaryotes (aerolysin-like proteins, ALPs) originated from recurrent horizontal gene transfer (HGT) (Moran et al., 2012). ALPs of different origins possess diverse functions, including immune defense and predation (Galinier et al., 2013; Szczesny et al., 2011; Xiang et al., 2014). The ALPs were significantly expanded in the genome of *C. heheva* (7 copies) compared with other echinoderms (0 or 1 copy) (*P* < 0.05) (**Table S10**). To investigate the possible origin and function of *C. heheva* ALPs, we reconstructed the phylogenetic tree of ALPs in echinoderms and diverse species. Interestingly, *C. heheva* ALPs did not cluster with ALPs from other echinoderms, suggesting that ALPs from *C. heheva* and other echinoderms have different origins (**Figure 6**). *Chiridota heheva* ALPs form a clade with ALPs from sea anemones (*Nematostella vectensis* and *Ecaiptasia diaphana*). This indicates that ALPs from *C. heheva* and sea anemones might have similar biological functions. It was shown that ALPs secreted by *N. vectensis* are involved in prey digestion (Moran et al., 2012). Therefore, the expansion of the ALP family in *C. heheva* might contribute to the disintegration of microbes during digestion.

**Figure 6.**
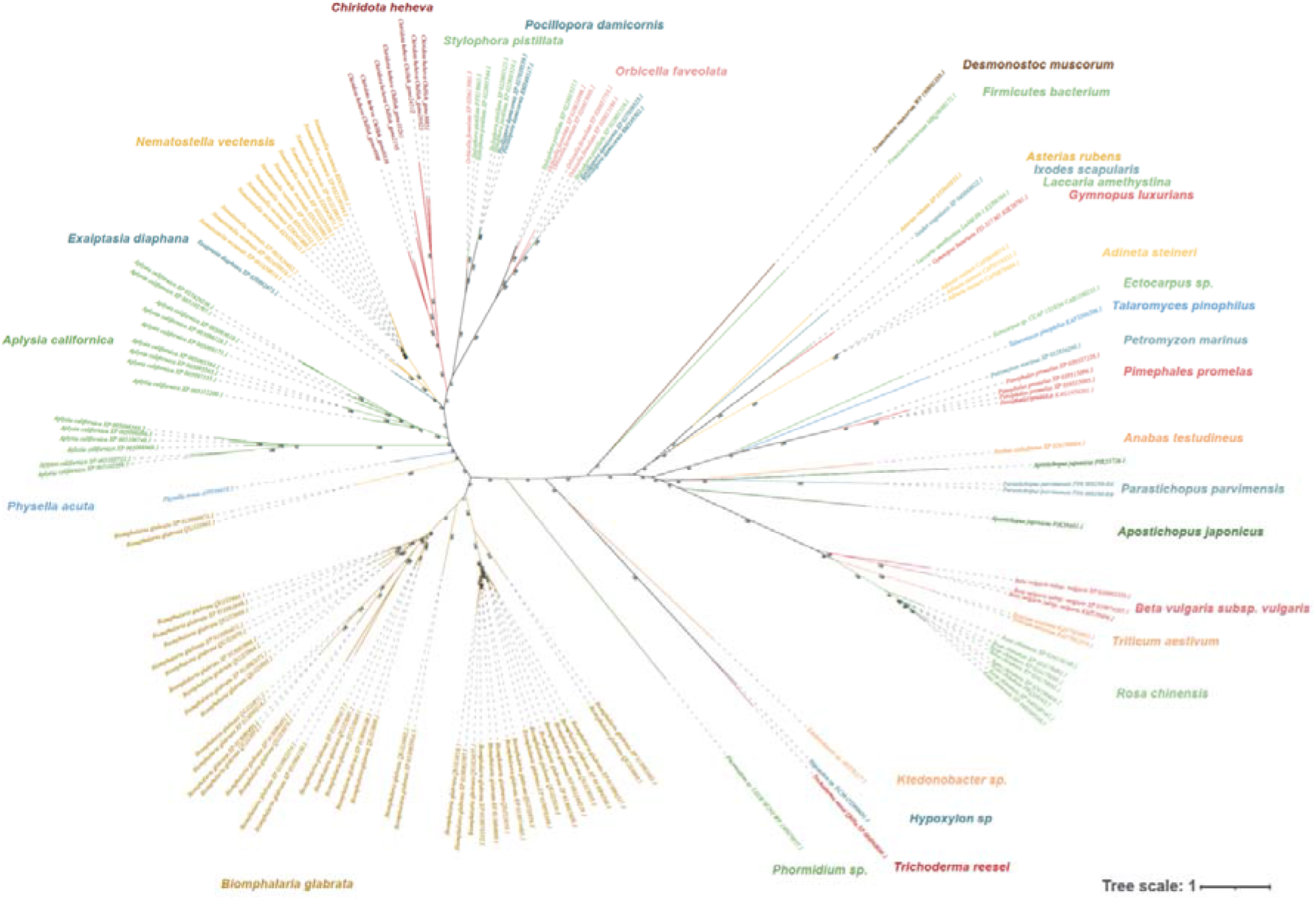
Evolutionary relationship with aerolysin-like proteins (ALPs) from *C. heheva* and other species. The unrooted phylogenetic tree was reconstructed using a maximum likelihood method. The values near the nodes are ultrafast bootstrap (UFBoot) values. ALPs from different types of species are highlighted by branches of different colors. The species name is shown near the corresponding lineage. ALPs from *C. heheva* do not cluster with ALPs from other echinoderms (*A. japonicus, P. parvimensis*), but with the ones from sea anemones (*N. vectensis, E. diaphana*).

### 3.7 Positively selected genes

To better understand the genetic basis of its adaptation to a deep-sea reducing environment, we identified genes undergoing positive selection (PSGs) in *C. heheva*. Compared with 6 other echinoderms, 27 PSGs were identified in the *C. heheva* genome (**Table S12**). Several hypoxia-related genes (*PKM, PAN2, LHPP*, and *RRP9*) (Benita et al., 2009; Bett et al., 2013; Chen et al., 2021; Luo et al., 2011), were subject to positive selection in *C. heheva*. Cold seeps and hydrothermal vents are characterized by low oxygen concentrations, which are challenging for endemic species (Hourdez & Lallier, 2007). Therefore, the adaptation of *C. heheva* to a deep-sea reducing environment may be attributed to selection against these hypoxia-related genes. Interestingly, the *LHPP* gene is also positively selected in the genomes of cetaceans, which are hypoxia-tolerant mammals (Tian et al., 2017). In addition, comparative genetic analysis showed that cetaceans and *C. heheva* have the same amino acid substitution at position 118 of the LHPP protein (**Figures S3 and S4**), which indicates a possible convergent evolution in the *LHPP* during the adaptation of cetaceans and *C. heheva* to hypoxic environments.

## 4. Discussion

With more than 1,400 extant species, Holothuroidea is one of the largest classes in the phylum Echinodermata (Pawson, 2007). In addition, holothurians are well adapted to diverse marine environments, with habitats ranging from shallow intertidal areas to hadal trenches (Jamieson, 2015; Smirnov et al., 2000). However, due to the lack of body fossils, evolutionary study of Holothuroidea is more difficult than other classes of Echinodermata. The high-quality genome of *C. heheva* presented in this report facilitates the investigation of its evolutionary history. Our phylogenomic analysis revealed that the divergence of echinoderms started in the early Cambrian (∼539 Ma), which is consistent with the fossil record. (Bottjer et al., 2006; Smith, 1988b) (**Fig. 3a**). The ancestor of *Chiridota heheva* diverged from the ancestors of two shallow-water holothurians (*A. japonicus* and *P. parvimensis*) approximately 429 Ma. As Apodida is the basal taxon in Holothuroidea, these results support the view that holothurians had evolved by the Early Ordovician (Reich, 2010). To better investigate the evolution of holothurians, we inferred the histories of ancestral population sizes of *C. heheva* and *A. japonicus* using PSMC (**Fig. 3b**). *Chiridota heheva* experienced a decline in population size approximately 21 Ma. Ocean temperature increased slowly between the late Oligocene and early Miocene (21-27 Ma) after long-term cooling from the end of the Eocene (Zachos et al., 1997; Zachos et al., 2001). Furthermore, species diversity within Echinodermata started to increase in the early Miocene (Kroh, 2007; Oyen & Portell, 2001). These results indicate that *C. heheva* might have colonized the current habitat in the early Miocene when the climate transition improved adaptations in echinoderms. The oceans experienced a decrease in temperature during the late Miocene (7 to 5.4 Ma) (Herbert et al., 2016). A decline in ancestral population size in *A. japonicus* started approximately 7 Ma. *Chiridota heheva* also experienced a moderate decline in population size in the early Pliocene. These results suggest that global cooling and environmental changes in the late Miocene were an important driver of demographic changes in both shallow-water and deep-sea holothurians.

Apodida do not have tube feet or complex respiratory trees, which are commonly found in other holothurians (Barnes, 1982). It was proposed that *Hox* genes might have contributed to the body development of echinoderms (Li et al., 2018). The gene composition and arrangement of the *Hox*/*ParaHox* gene cluster were consistent between *A. japonicus* and *C. heheva*, indicating that *Hox* genes were unlikely to have been involved in the morphological divergence between Apodids and other holothurians. There are some inconsistent results regarding the gene composition of *Hox* gene clusters in different echinoderm genomes. Previous studies found that *Hox4* and *Hox6* were missing in the genomes of holothurians (Li et al., 2018; Zhang et al., 2017), and *Hox4* was missing in the genomes of echinoids (Cameron et al., 2006). Li *et al*. (2018) proposed that *Hox6* was lost before the split of Echinozoa and Asterozoa (Li et al., 2018), while Li *et al*. (2020) suggested that the loss of *Hox4* or *Hox6* was a lineage-specific event (Li et al., 2020). We found that *Hox4* and *Hox 6* were missing in the genomes of both *C. heheva* and *A. japonicus*. In addition, *Hox6* was missing in the genome of *A. planci*, and *Hox4* was missing in the genomes of both *L. variegatus* and *S. purpuratus* (**Fig. 4a**). This suggests that *Hox4* was lost before the split of Echinoidea and Holothuroidea, and *Hox6* was lost in Holothuroidea and Asteroidea. This scenario is not parsimonious, as Holothuroidea is paraphyletic with Asteroidea. As *S. purpuratus Hox6* clusters with *A. planci Hox4* in phylogenetic analysis, Baughman et al. (2014) proposed reclassifying *S. purpuratus Hox6* as *Hox4* (Baughman et al., 2014). Following this argument, *Hox6* was missing in Holothuroidea, Echinoidea, and Asteroidea, and *Hox4* was missing in Holothuroidea (**Fig. 4b**). This supports the view that the loss of *Hox6* occurred before the split of Echinozoa and Asterozoa.

Comparative genomic analysis showed that the ALP gene family was significantly expanded in *C. heheva* compared with other echinoderms. The expansion of the ALP family in *C. heheva* might have contributed to its adaptation to cold seep environments. Cold seeps are areas where methane, hydrogen sulfide, and other hydrocarbons seep or emanate as gas from deep geologic sources (Suess, 2014). Chemosynthetic microbes oxidize the reduced chemicals contained in the fluids to produce energy and fix carbon into organic matter, which supports large benthic communities around the gas source (Levin, 2005). Most seep-dwelling animals survive by hosting chemosynthetic microbes (Petersen & Dubilier, 2009). *Chiridota heheva* has a unique feeding habit of acquiring nutrients from sediment detritus, suspended material, and wood fragments when available. The microbial communities of cold seeps are very different from those of other seafloor ecosystems (Ruff et al., 2015). Moreover, some of these microbes have unique cellular structures that might be difficult to disintegrate (Katayama et al., 2020), which impedes nutrient acquisition of *C. heheva* from free-living microbes of cold seeps. As typical pore-forming proteins, aerolysin and related proteins are found in a large variety of species and possess diverse functions (Szczesny et al., 2011). It was proposed that ALPs were derived from recurrent horizontal gene transfer. ALPs of the same origin might have similar functions (Moran et al., 2012). *Chiridota heheva* ALPs and ALPs from other echinoderms are likely to have different origins, as they were clustered with aerolysins from distinct groups of bacteria (**Figure 6**). *Chiridota heheva* ALPs formed a clade with sea anemone ALPs. Furthermore, ALPs from hydra and sea anemones are involved in prey disintegration after predation by lysing cells through pore formation on membranes (Moran et al., 2012; Sher et al., 2008). This suggests that the expansion of the ALP family might have contributed to microbe digestion in *C. heheva*, which in turn facilitated its adaptation to cold seep environments.

Several genes that are involved in hypoxic responses (*PKM, PAN2*, and *LHPP*) and one of the HIF-1 target genes (*PPR9*) were subjected to positive selection in *C. heheva*. The transcription of the *PKM2* gene is activated by HIF-1. PKM2 promotes transactivation of HIF-1 target genes by directly interacting with the HIF-1α subunit. PKM2 is involved in a feedback loop that reprograms glucose metabolism under hypoxic conditions (Luo et al., 2011). LHPP induces ubiquitin-mediated degradation of PKM2, which results in the inhibition of glycolysis under hypoxia (Chen et al., 2021). Interestingly, the LHPP was also subject to positive selection in cetaceans (Tian et al., 2017). Furthermore, both *C. heheva* and cetaceans have the same amino acid substitution at position 118 of the LHPP protein (**Figs. S3 and S4**). These results suggest that the two interacting genes (*PKM2* and *LHPP*) play a key role in the hypoxic adaptation of these hypoxia-tolerant marine animals.

## Acknowledgements

We thank Dr. Kang Ding, and Dr. Zhimin Jian for leading the expedition of TS12-02, the crew of research vessel (R/V) *Tansuoyihao*, the pilot team of the manned submersible *Shenhaiyongshi*, and the onboard diving scientists for their technical support during the cruise. We gratefully acknowledge the National Supercomputing Center in Guangzhou for provision of computational resources. This study was supported by Innovation Group Project of Southern Marine Science and Engineering Guangdong Laboratory (Zhuhai) (No. 311021006), National Natural Science Foundation of China (No. 31900309), GuangDong Basic and Applied Basic Research Foundation (No. 2019A1515011644), and National Innovation and Entrepreneurship Training Project for College Student of China (No. 20201126).

## Data Accessibility

Raw reads and genome assembly are accessible in NCBI under BioProject number PRJNA752986. Assembled genome sequences are accessible under Whole Genome Shotgun project number JAIGNY000000000. Raw reads and genome assembly are also available at the CNGB Sequence Archive (CNSA) of China National GeneBank DataBase (CNGBdb) with accession number CNP0002134. The genome assembly and related annotation files are available at Figshare (https://doi.org/10.6084/m9.figshare.15302229).

## Author Contributions

M.W and J.G.H. conceived of the project and designed research; J.H. collected the sample; P.T., L.Z, Y.M., Q.C., Q.Z., L.Z. assembled and annotated the genome; L.Z., Z.G., J.H., M.W., S.Q., Y.W. performed the evolutionary analyses; M.W., G.H. wrote the paper the manuscript with the contribution from all authors.

## Notes

### Competing Interest Statement

The authors have declared no competing interest.

