## Supplementary Materials for "The genome of an apodid holothuroid (*Chiridota heheva*) provides insights into its adaptation to deep-sea reducing environment"

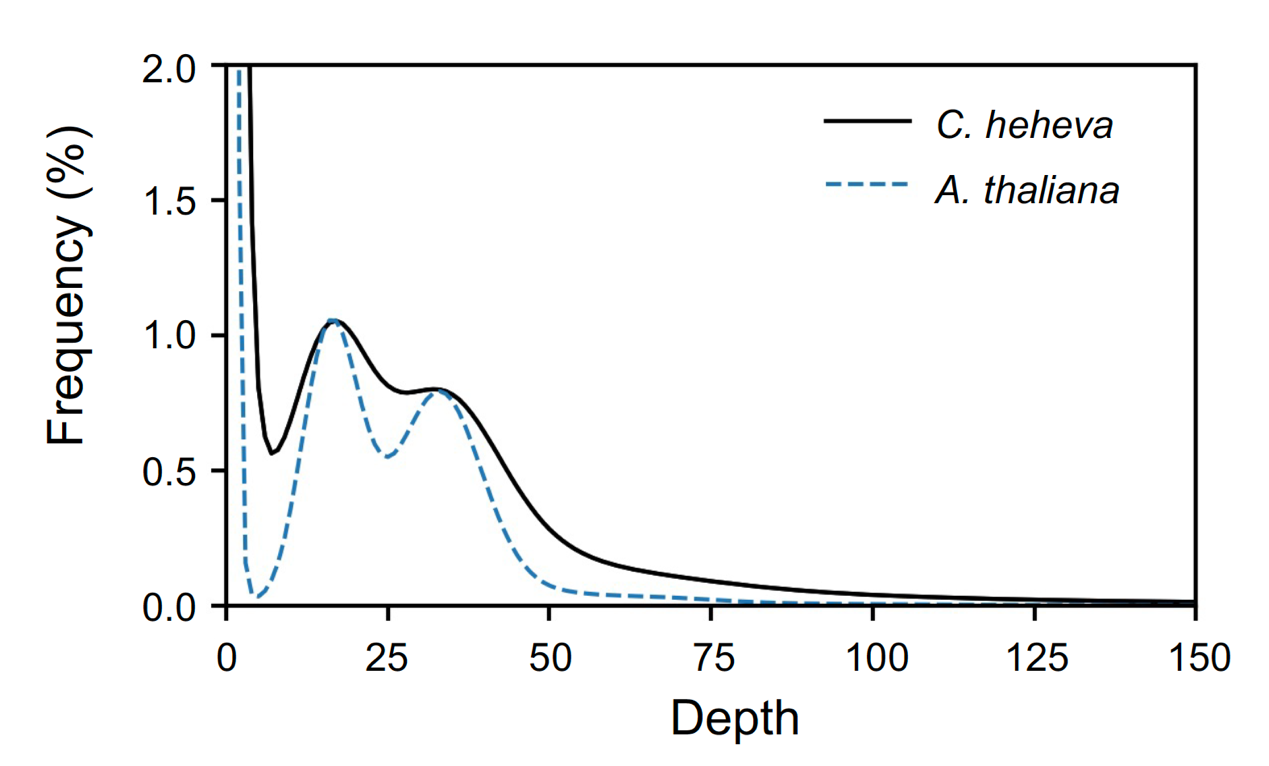

**Figure S1. Distribution of 17-mer frequency in *C. heheva* genome.** The heterozygous rate and the genome size were determined based on the *k*-mer distribution. The average coverage depth is estimated to be 34 for *C. heheva* based on 150 bp paired-end Illumina reads**.**

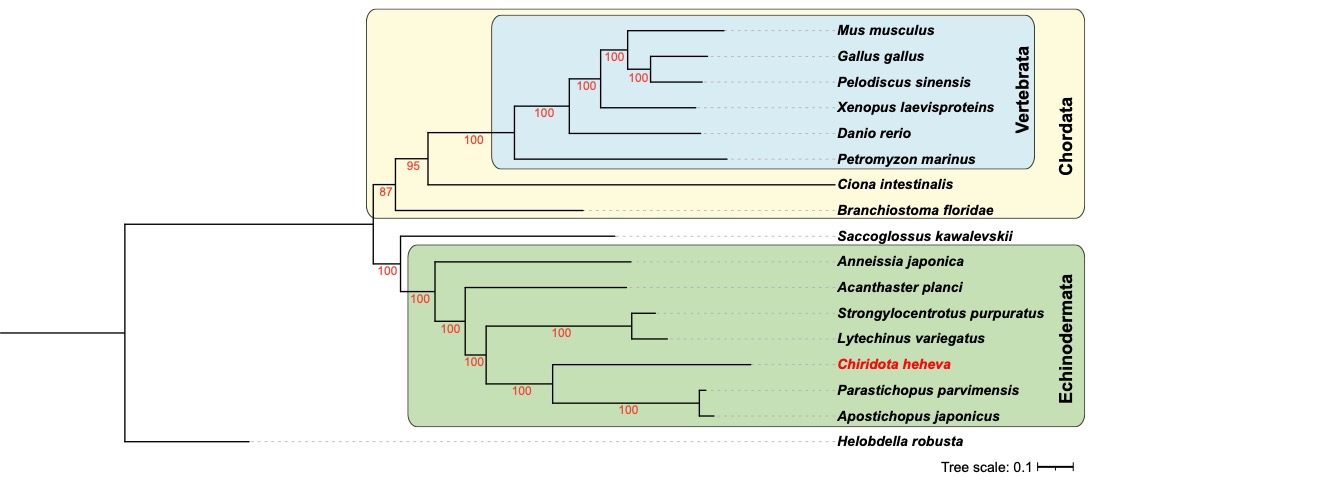

**Figure S2. The phylogenetic tree of *C. heheva* and 16 other metazoans.** The tree was reconstructed with 80 single-copy orthologs using a maximum likelihood approach. The ultrafast bootstrap (UFBoots) value is listed below each of the nodes.

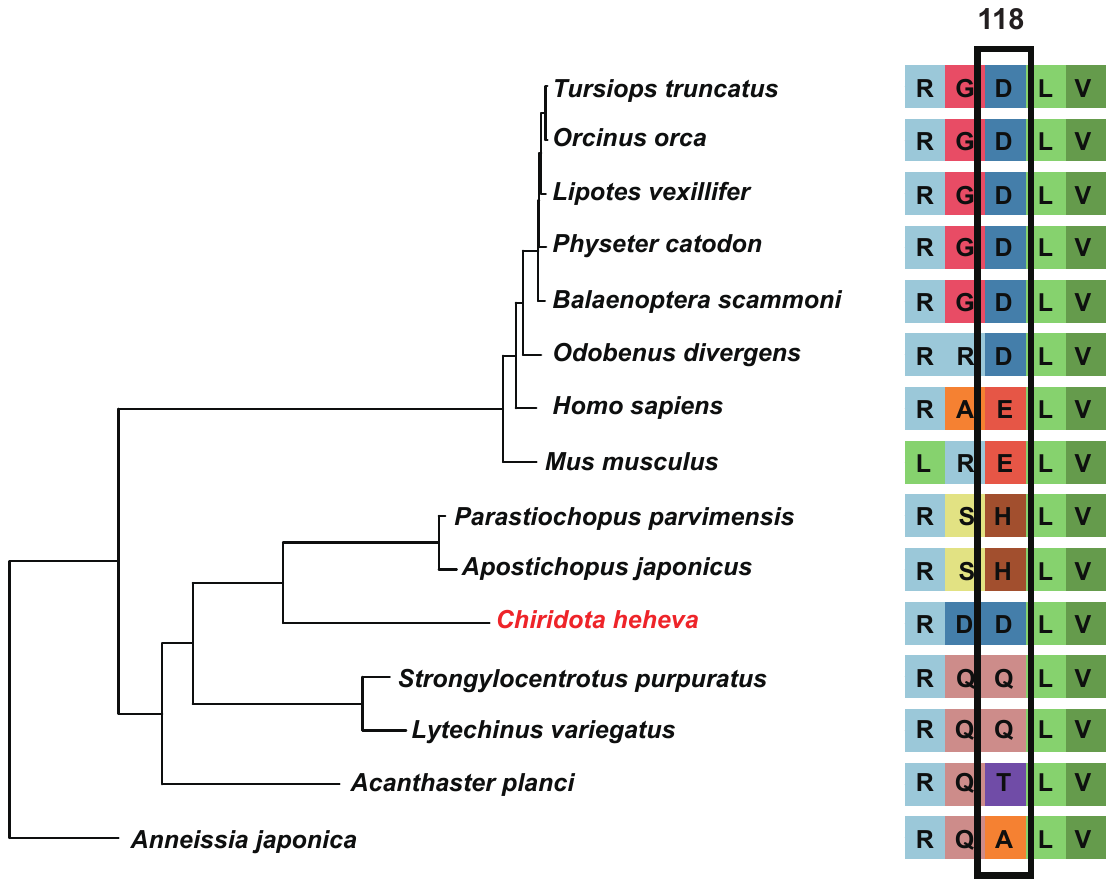

**Figure S3. A possible amino acid substitution of LHPP that contributed to hypoxic adaptation in *C. heheva* and cetaceans.** The maximum likelihood phylogenetic tree of cetaceans, *C. heheva* and other echinoderms was reconstructed using 598 single-copy orthologs. *C. heheva* and cetaceans, which are tolerant to hypoxia, have the same amino acid substitution at position 118 of LHPP protein.

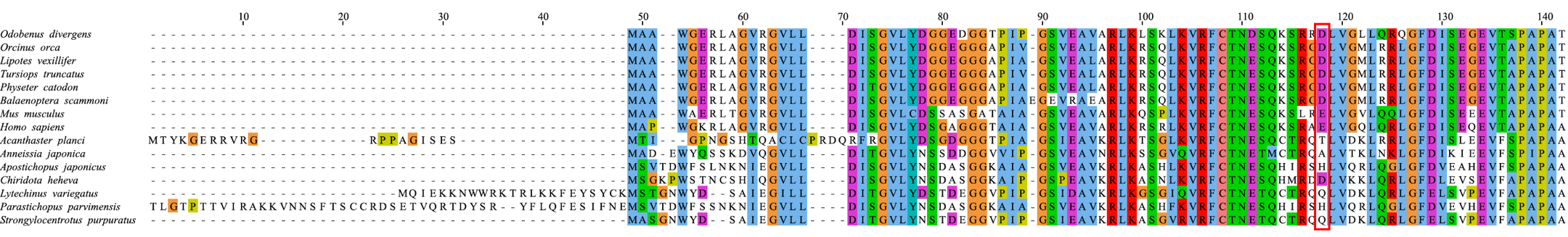

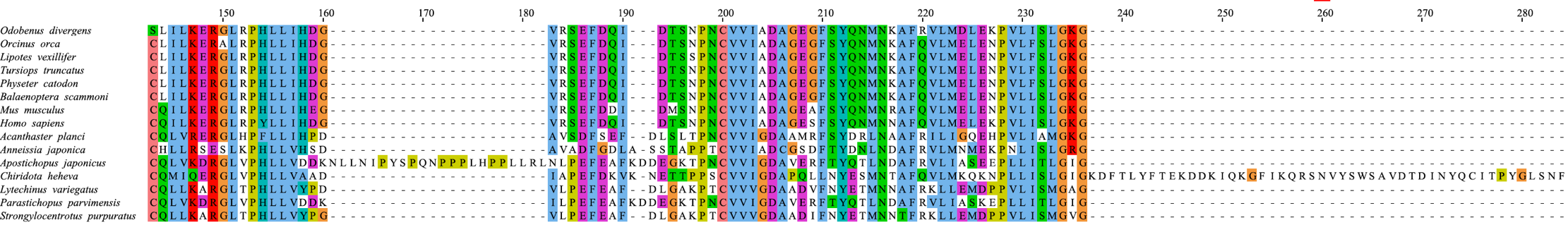

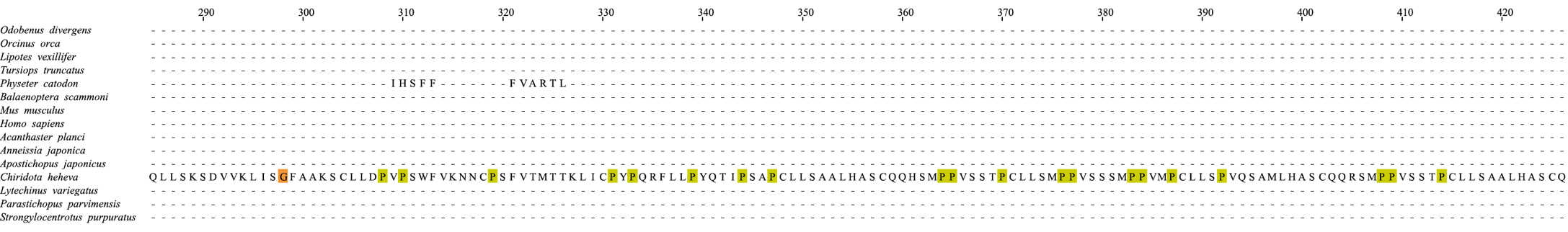

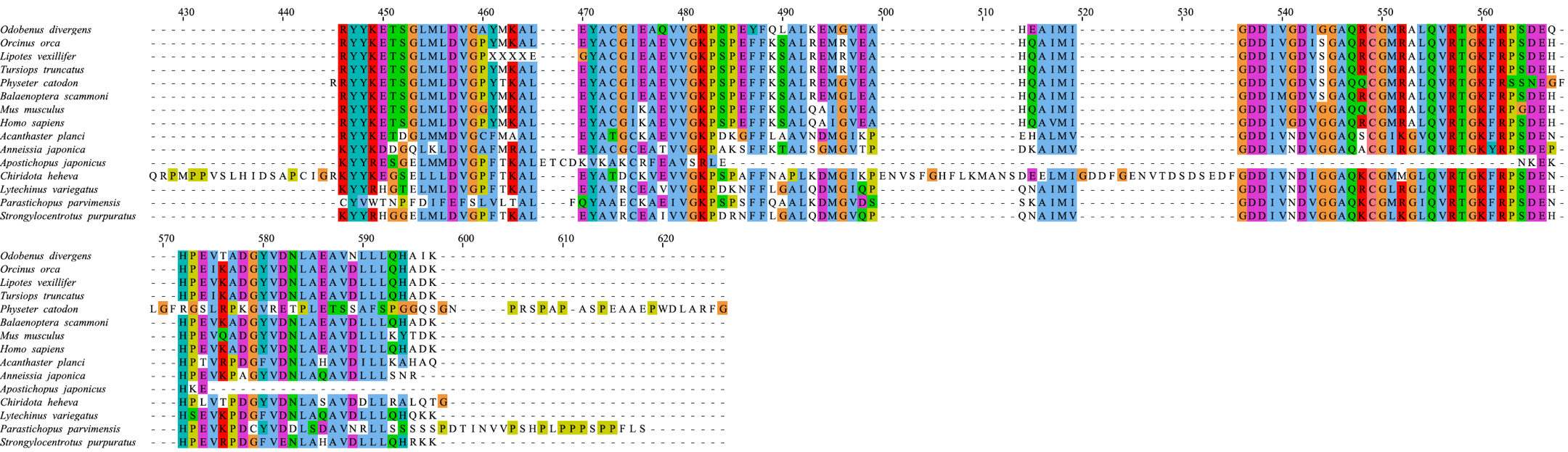

**Figure S4. Multi-sequence alignment of LHPP from cetaceans, *C. heheva*, and other echinoderms.** *C. heheva* and cetacean have a same amino acid substitution at position 118 of LHPP.

**Table S1 Basic statistics of Nanopore reads**

| **Total number of reads** | **Total number of bases (Gb)** | **Maximum length of reads (bp)** | **Minimum length of reads (bp)** | **Average length of reads (bp)** | **Depth of coverage** |
| --- | --- | --- | --- | --- | --- |
| 2,905,304 | 42.43 | 246,442 | 25 | 1460.4 | 34 |

**Table S2 Basic statistics of Illumina reads**

| **Total number of reads** | **Total number of base pairs** | **Percentage of Q20 base pairs (%)** | **Percentage of Q30 base pairs (%)** | **GC (%)** |
| --- | --- | --- | --- | --- |
| 357,304,954 | 49,193,650,851 | 96.4 | 90.83 | 38.38 |

**Table S3 Summary of *k*-mer analysis**

| ***k*-mer** | ***k*-mer**  **number** | **k-mer depth** | **Genome size (bp)** | **Heterozygosity (%)** |
| --- | --- | --- | --- | --- |
| 17 | 41,978,881,806 | 34 | 1,234,672,994 | 2.0 |

**Table S4 BUSCO evaluation of *C. heheva* genome assembly**

|  | *C. heheva* |
| --- | --- |
| Complete BUSCOs | 855 |
| Complete and single-copy BUSCOs | 849 |
| Complete and duplicated BUSCOs | 6 |
| Fragmented BUSCOs | 27 |
| Missing BUSCOs | 72 |
| Total BUSCO groups searched | 954 |

**Table S5 Summary of SQUAT analysis**

|  | **Statistics** |
| --- | --- |
| No. of sequence | 332,594,866 |
| Sample size | 1,000,000 |
| Sequence length | 15-140 |
| Avgerage poorly mapped sequence | 8.9% |
| GC content | 38% |

**Table S6 Summary of annotated repeats in *C. heheva* genome**

|  | Number | Length（bp） | Percentage（%） |
| --- | --- | --- | --- |
| **Retroelements** | **277,203** | **126,998,333** | **11.47** |
| SINEs: | 43,231 | 7,207,532 | 0.65 |
| Penelope | 9,627 | 3,239,963 | 0.29 |
| LINEs: | 210,096 | 107,581,691 | 9.72 |
| L2/CR1/Rex | 96,117 | 41,383,441 | 3.74 |
| R1/LOA/Jockey | 63,290 | 44,118,706 | 3.99 |
| R2/R4/NeSL | 125 | 64,396 | 0.01 |
| RTE/Bov-B | 28,298 | 11,944,523 | 1.08 |
| L1/CIN4 | 418 | 66,470 | 0.01 |
| LTR: | 23,876 | 12,209,110 | 1.1 |
| BEL/Pao | 4,654 | 792,365 | 0.07 |
| Gypsy/DIRS1 | 17,877 | 11,065,063 | 1 |
| Retroviral | 1,294 | 295,882 | 0.03 |
| **DNA transposons：** | **79,045** | **33,585,526** | **3.03** |
| hobo-Activator | 36,285 | 9,130,721 | 0.82 |
| Tc1-IS630-Pogo | 849 | 286,365 | 0.03 |
| PiggyBac | 158 | 81,861 | 0.01 |
| Tourist/Harbinger | 970 | 346,863 | 0.03 |
| Other (Mirage,P-element,Transib) | 3,621 | 539,664 | 0.05 |
| **Rolling circles** | **34,565** | **9,570,594** | **0.86** |
| **Unclassified:** | **1,745,539** | **424,910,357** | **38.39** |
| **Small RNA:** | **7,394** | **1,286,798** | **0.12** |
| **Satellites:** | **6,112** | **1,122,725** | **0.1** |
| **Simple repeats:** | **52,606** | **26,896,646** | **2.43** |

**Table S7** **Transposable element composition in *C. heheva* and other echinoderm genomes**

|  | *Chiridota*  *heheva*  (1,107 Mb) | | | *Apostichopus japonicus*  (952 Mb) | | | *Parastichopus parvimensis*  (873 Mb) | | | *Strongylocentrotus*  *purpuratus*  (991 Mb) | | | *Lytechinus variegatus*  (1,061 Mb) | | | *Acanthaster*  *planci*  (384Mb) | | | *Anneissia*  *Japonica*  (553 Mb) | |
| --- | --- | --- | --- | --- | --- | --- | --- | --- | --- | --- | --- | --- | --- | --- | --- | --- | --- | --- | --- | --- |
|  | Length (Mb) | Rate (%) | Length (Mb) | | Rate (%) | Length (Mb) | | Rate (%) | Length (Mb) | | Rate (%) | Length (Mb) | | Rate (%) | Length (Mb) | | Rate (%) | Length (Mb) | | Rate (%) |
| DNA | 33.59 | 3.03 | 34.56 | | 3.52 | 10.31 | | 1.18 | 105.74 | | 10.67 | 211.9 | | 19.97 | 40.12 | | 10.45 | 9.56 | | 1.62 |
| LTR | 12.21 | 1.10 | 9.63 | | 0.98 | 2.71 | | 0.31 | 12.04 | | 1.22 | 32.56 | | 3.07 | 14.82 | | 3.86 | 5.62 | | 0.95 |
| LINE | 107.58 | 9.72 | 20.33 | | 2.07 | 15.57 | | 1.78 | 37.88 | | 3.82 | 54.94 | | 5.18 | 18.28 | | 4.76 | 27.16 | | 4.61 |
| SINE | 7.21 | 0.65 | 5.89 | | 0.60 | 2.57 | | 0.29 | 15.12 | | 1.53 | 32.4 | | 3.05 | 4.22 | | 1.10 | 6.07 | | 1.03 |
| RNA | 1.29 | 0.12 | 0.76 | | 0.08 | 3.25 | | 0.4 | 0.16 | | 0.02 | 0.15 | | 0.01 | 26.17 | | 6.82 | 1.51 | | 0.26 |
| Tandem Repeat | 28.02 | 2.53 | 19.34 | | 1.97 | 1.34 | | 0.16 | 40.23 | | 4.06 | 25.31 | | 2.39 | 3.41 | | 0.89 | 1.95 | | 0.33 |
| Other | 9.57 | 0.86 | 12.21 | | 1.24 | 0.58 | | 0.07 | 5.74 | | 0.58 | 48.61 | | 4.58 | 13.61 | | 3.54 | 1.08 | | 0.18 |
| Unclassified | 424.91 | 38.39 | 151.26 | | 15.39 | 181.87 | | 20.83 | 135.2 | | 13.64 | 18.69 | | 0.02 | 5.17 | | 1.35 | 215.6 | | 36.56 |
| Total | 624.38 | 56.40 | 253.98 | | 26.68 | 218.2 | | 25.02 | 352.11 | | 35.54 | 424.56 | | 38.27 | 125.8 | | 32.77 | 268.55 | | 45.54 |

**Table S8 Gene features of *C. heheva* and other echinoderms**

|  | Gene counts | Average gene length (bp) | Average CDS length (bp) | Average exons per gene | Average exon size (bp) | Average intron size (bp) |
| --- | --- | --- | --- | --- | --- | --- |
| *C. heheva* | 36,527 | 14,943 | 1,496 | 5.37 | 277 | 2,859 |
| *A. japonicus* | 29,451 | 7,722 | 1,324 | 7.40 | 202 | 1,134 |
| *S. purpuratus* | 27,750 | 13,690 | 2,143 | 8.90 | 282 | 1,217 |
| *L. variegatus* | 28,094 | 18,033 | 1,062 | 5.30 | 198 | 1,527 |
| *A. planci* | 24,747 | 16,844 | 1,375 | 6.80 | 203 | 1,161 |

**Table S9 Summary of NLR genes in *C. heheva***

| **NLR** | **Domain organization** |
| --- | --- |
| ChiHeh_gene29120 | Death, NACHT, Death |
| ChiHeh_gene12421 | DED, NACHT |
| ChiHeh_gene26187 | NACHT |
| ChiHeh_gene10961 | DED, DED, NACHT |
| ChiHeh_gene15903 | NACHT |
| ChiHeh_gene26414 | DED, NACHT |
| ChiHeh_gene10067 | DED, DED, NACHT |
| ChiHeh_gene35009 | DED, DED, NACHT |
| ChiHeh_gene30081 | Death, NACHT |
| ChiHeh_gene12883 | DED, NACHT |
| ChiHeh_gene25938 | Death, NACHT |
| ChiHeh_gene1054 | NACHT |
| ChiHeh_gene10777 | Death, NACHT |
| ChiHeh_gene19441 | NACHT |
| ChiHeh_gene28117 | DED, DED, NACHT |
| ChiHeh_gene10117 | NACHT |
| ChiHeh_gene31453 | DED, DED, NACHT |
| ChiHeh_gene10309 | DED, DED, NACHT |
| ChiHeh_gene34531 | Death, NACHT |
| ChiHeh_gene14426 | DED, NACHT |
| ChiHeh_gene8030 | NACHT |
| ChiHeh_gene12018 | NACHT, LRR_8, LRR_8, LRR_4 |
| ChiHeh_gene4819 | Death, NACHT, DDE_Tnp_1_7 |
| ChiHeh_gene20564 | NACHT |
| ChiHeh_gene3835 | NACHT |
| ChiHeh_gene3488 | NACHT |
| ChiHeh_gene7016 | NACHT |
| ChiHeh_gene3262 | DED, DED, NACHT |
| ChiHeh_gene7544 | NACHT |
| ChiHeh_gene7545 | DED, NACHT |
| ChiHeh_gene26021 | DED, DED, NACHT |
| ChiHeh_gene10245 | V-set, V-set, C2-set_2, NACHT |
| ChiHeh_gene12145 | NACHT |
| ChiHeh_gene11999 | NACHT |
| ChiHeh_gene30189 | Death, NACHT |
| ChiHeh_gene33098 | NACHT |
| ChiHeh_gene2291 | V-set, Ig_3, NACHT |
| ChiHeh_gene32597 | NACHT, ZU5 |
| ChiHeh_gene17853 | DED, NACHT |
| ChiHeh_gene25299 | NACHT |
| ChiHeh_gene25442 | NACHT |
| ChiHeh_gene25963 | V-set, C2-set_2, NACHT |
| ChiHeh_gene29023 | DED, NACHT, DED |
| ChiHeh_gene7279 | NACHT |
| ChiHeh_gene18722 | NACHT |
| ChiHeh_gene1587 | NACHT |
| ChiHeh_gene7919 | NACHT |
| ChiHeh_gene23708 | NACHT |
| ChiHeh_gene7425 | NACHT |
| ChiHeh_gene16374 | DED, DED, NACHT |
| ChiHeh_gene19652 | Death, NACHT |
| ChiHeh_gene3439 | V-set, V-set, C2-set_2, NACHT, zf-B_box, zf-B_box |
| ChiHeh_gene22493 | NACHT, NLRC4_HD2 |

**Table S10 Gene families that are expanded in *C. heheva* compared to other echinoderms**

| Pather ID | Annotation | C. heheva | A. japonicus | P. parvimensis | S. purpuratus | L. variegatus | A. planci | A. japonica |
| --- | --- | --- | --- | --- | --- | --- | --- | --- |
| PTHR22984 | Serine/Threonine-Protein Kinase PIM | 67 | 0 | 0 | 1 | 1 | 1 | 1 |
| PTHR35365 | LP04239P | 60 | 0 | 0 | 0 | 0 | 0 | 1 |
| PTHR13582 | M-phase phosphoprotein 6 | 48 | 1 | 1 | 1 | 1 | 1 | 1 |
| PTHR36493 | unknown | 45 | 0 | 0 | 0 | 0 | 0 | 1 |
| PTHR18847 | 20 kD Nuclear Cap Binding Protein | 40 | 2 | 0 | 1 | 1 | 1 | 1 |
| PTHR14136 | Uncharacterized | 38 | 0 | 0 | 0 | 0 | 0 | 1 |
| PTHR35577 | Cysteine-Rich, Acidic Integral Membrane Protein-Related | 37 | 0 | 0 | 0 | 0 | 0 | 1 |
| PTHR12366 | Aspartyl/Asparaginyl β-Hydroxylase | 32 | 2 | 0 | 1 | 1 | 1 | 1 |
| PTHR36910 | Unknown | 28 | 0 | 0 | 0 | 0 | 0 | 1 |
| PTHR23080 | THAP-type zinc finger | 27 | 1 | 2 | 3 | 2 | 1 | 3 |
| PTHR36144 | S-Antigen Protein | 26 | 1 | 0 | 0 | 0 | 1 | 1 |
| PTHR12220 | 50S/60S Ribosomal Protein L16 | 25 | 2 | 0 | 2 | 1 | 1 | 1 |
| PTHR46670 | Unknown | 25 | 1 | 0 | 1 | 1 | 1 | 1 |
| PTHR13160 | Oligosaccharyl transferase complex, subunit OST3/OST6 | 18 | 2 | 0 | 1 | 1 | 1 | 1 |
| PTHR33480 | Unknown | 16 | 1 | 0 | 2 | 1 | 0 | 2 |
| PTHR46880 | Unknown | 14 | 0 | 2 | 0 | 0 | 0 | 2 |
| PTHR15135 | STAC | 13 | 1 | 1 | 0 | 0 | 0 | 1 |
| PTHR19143 | Fibrinogen, alpha/beta/gamma chain, C-terminal globular | 12 | 3 | 1 | 3 | 4 | 4 | 2 |
| PTHR11239 | DNA-Directed RNA Polymerase | 12 | 1 | 2 | 1 | 1 | 1 | 1 |
| PTHR46585 | Integrase Core Domain Containing Protein | 12 | 1 | 0 | 0 | 0 | 3 | 3 |
| PTHR47883 | Unknown | 12 | 0 | 0 | 0 | 0 | 0 | 1 |
| PTHR13681 | Survival Of Motor Neuron-Related-Splicing Factor 30-Related | 11 | 1 | 1 | 2 | 1 | 1 | 2 |
| PTHR14859 | Calcofluor White Hypersensitive Protein Precursor | 11 | 0 | 2 | 1 | 1 | 1 | 1 |
| PTHR48039 | RNA-Binding Motif Protein 14B | 10 | 1 | 2 | 2 | 2 | 2 | 2 |
| PTHR10408 | Sterol O-acyltransferase, ACAT/DAG/ARE types | 10 | 1 | 0 | 1 | 1 | 1 | 1 |
| PTHR47202 | Uncharacterized | 10 | 0 | 0 | 0 | 1 | 0 | 1 |
| PTHR24559 | Transposon Ty3-I Gag-Pol Polyprotein | 9 | 0 | 0 | 1 | 0 | 0 | 1 |
| PTHR23184 | Tetratricopeptide Repeat Protein 14 | 8 | 3 | 1 | 2 | 1 | 1 | 1 |
| PTHR11584 | Serine/Threonine Protein Kinase | 8 | 1 | 1 | 1 | 2 | 1 | 1 |
| PTHR46221 | FERM and PDZ Domain-Containing Protein Family Member | 8 | 0 | 2 | 0 | 0 | 0 | 1 |
| PTHR13415 | Integrator Complex Subunit | 7 | 1 | 1 | 2 | 1 | 1 | 1 |
| PTHR34007 | Aerolysin-like Protein-Related | 7 | 1 | 0 | 1 | 2 | 1 | 1 |
| PTHR24134 | Ankyrin repeat-containing domain superfamily | 7 | 2 | 1 | 0 | 0 | 0 | 0 |
| PTHR13105 | Myeloid Leukemia Factor 11 | 7 | 1 | 0 | 1 | 1 | 1 | 1 |
| PTHR46670 | Uncharacterized | 7 | 1 | 0 | 1 | 1 | 0 | 1 |
| PTHR12247 | Polycomb Group Protein | 6 | 1 | 2 | 1 | 2 | 1 | 1 |
| PTHR12867 | Glycosyl Transferase-Related | 6 | 1 | 0 | 1 | 1 | 1 | 1 |
| PTHR47522 | Salvador Family WW Domain-Containing Protein 1 | 6 | 1 | 0 | 1 | 1 | 1 | 1 |
| PTHR11506 | Lysosome-Associated Membrane Glycoprotein | 6 | 1 | 0 | 1 | 0 | 1 | 1 |
| PTHR18935 | Uncharacterized | 6 | 0 | 0 | 1 | 1 | 0 | 1 |
| PTHR23184 | Tetratricopeptide Repeat Protein 14 | 6 | 0 | 0 | 0 | 0 | 1 | 1 |
| PTHR24559 | Transposon Ty3-I Gag-Pol Polyprotein | 6 | 0 | 0 | 0 | 0 | 0 | 1 |
| PTHR46599 | Piggybac Transposable Element-Derived Protein | 6 | 0 | 0 | 0 | 0 | 0 | 1 |
| PTHR11929 | Alpha-1,3-Fucosyltransferase | 5 | 1 | 1 | 6 | 6 | 0 | 1 |
| PTHR21567 | CLASP | 5 | 2 | 1 | 1 | 2 | 2 | 1 |
| PTHR21368 | 50S Ribosomal Protein L9 | 5 | 1 | 2 | 1 | 2 | 1 | 1 |
| PTHR20938 | Uncharacterized | 5 | 1 | 0 | 2 | 2 | 1 | 1 |
| PTHR22890:SF2 | Mediator Of RNA Polymerase II Transcription Subunit 11 | 5 | 1 | 0 | 2 | 1 | 1 | 1 |
| PTHR46495 | Dual Specificity Protein Phosphatase | 5 | 1 | 0 | 2 | 1 | 1 | 1 |
| PTHR19315 | ER Membrane Protein Complex Subunit | 5 | 1 | 0 | 1 | 1 | 1 | 1 |
| PTHR46079 | FERM Domain-Containing Protein 4 | 5 | 1 | 0 | 0 | 0 | 0 | 1 |
| N/A | No Annotation | 5 | 1 | 0 | 0 | 0 | 0 | 1 |
| PTHR11830:SF3 | Cylindromatosis, Isoform D | 5 | 1 | 0 | 0 | 0 | 0 | 1 |
| PTHR24240 | OPSIN | 4 | 2 | 0 | 2 | 2 | 1 | 2 |
| PTHR46954 | Uncharacterized | 4 | 2 | 0 | 1 | 2 | 1 | 2 |
| PTHR14628 | BEN Domain-Containing Protein 5 | 4 | 2 | 0 | 1 | 2 | 1 | 1 |
| PTHR14338 | Actin Filament-Associated Protein 1 Family Member | 4 | 1 | 0 | 1 | 2 | 1 | 1 |
| PTHR19303 | Transposon | 4 | 0 | 1 | 0 | 1 | 1 | 1 |
| PTHR45036 | Methyltransferase Like 7B | 3 | 2 | 0 | 1 | 2 | 2 | 2 |
| PTHR11575 | 5'-Nucleotidase-Related | 3 | 2 | 0 | 2 | 2 | 1 | 2 |
| PTHR46007 | Mediator Of RNA Polymerase II Transcription Subunit 12 | 3 | 2 | 0 | 2 | 1 | 1 | 2 |
| PTHR45641 | Tetratricopeptide Repeat Protein | 3 | 2 | 0 | 2 | 2 | 0 | 2 |
| PTHR33442 | Trans-3-Hydroxy-L-Proline Dehydratase | 3 | 2 | 0 | 1 | 2 | 1 | 1 |
| PTHR31664 | Protein CBG16427 | 3 | 2 | 0 | 0 | 0 | 2 | 1 |
| PTHR11347 | Cyclic Nucleotide Phosphodiesterase | 3 | 1 | 0 | 0 | 0 | 0 | 1 |
| PTHR47018 | Uncharacterized | 2 | 1 | 0 | 0 | 1 | 0 | 1 |

**Table S11 Gene families that are contracted in *C. heheva* compared to other echinoderms**

| ***PANTHER ID*** | ***Annotation*** | **C. heheva** | **A. japonicus** | **P. parvimensis** | **S. purpuratus** | **L. variegatus** | **A. planci** | **A. japonica** |
| --- | --- | --- | --- | --- | --- | --- | --- | --- |
| *PTHR42743* | Amino-Acid Aminotransferase | 2 | 2 | 2 | 1 | 1 | 13 | 16 |
| *PTHR14002* | Endoglin/Tgf-β Receptor Type III | 1 | 2 | 2 | 1 | 1 | 5 | 23 |
| *PTHR14948* | NG5 | 1 | 3 | 5 | 1 | 1 | 2 | 11 |
| *PTHR10578* | S-2-Hydroxy-Acid Oxidase-Related | 1 | 2 | 4 | 3 | 2 | 2 | 3 |
| *PTHR11733* | Zinc Metalloprotease Family M13 Neprilysin-Related | 0 | 1 | 1 | 1 | 1 | 11 | 2 |
| *PTHR22930* | Uncharacterized | 0 | 1 | 2 | 0 | 0 | 1 | 10 |
| *PTHR10974* | Uncharacterized | 0 | 1 | 2 | 2 | 1 | 1 | 6 |
| *PTHR43586* | Cysteine Desulfurase | 0 | 2 | 1 | 2 | 4 | 1 | 2 |
| *PTHR11841* | Reelin | 0 | 4 | 2 | 2 | 1 | 1 | 2 |
| *PTHR47968* | Centromere Protein E | 0 | 5 | 2 | 1 | 1 | 1 | 1 |
| *PTHR23033* | β-1,3-Galactosyltransferase | 0 | 1 | 2 | 4 | 1 | 1 | 1 |
| *PTHR12555* | Ubiquitin Fusion Degradation Protein | 0 | 3 | 1 | 1 | 1 | 3 | 1 |
| *PTHR14237* | Molybdopterin Cofactor Sulfurase Mosc | 0 | 1 | 4 | 2 | 1 | 1 | 1 |
| *PTHR14633* | Little Elongation Complex Subunit 2 | 0 | 2 | 1 | 3 | 1 | 2 | 1 |
| *PTHR22367* | Coiled-Coil Domain-Containing Protein 14 | 0 | 2 | 1 | 3 | 1 | 1 | 1 |
| *PTHR11730* | Ammonium Transporter | 0 | 3 | 1 | 2 | 1 | 1 | 1 |
| *PTHR19981* | Talin | 0 | 4 | 1 | 1 | 1 | 1 | 1 |
| *PTHR28434* | Protein C3ORF33 | 0 | 1 | 2 | 3 | 1 | 1 | 1 |
| *PTHR31804* | Mediator Of RNA Polymerase Ii Transcription Subunit | 0 | 3 | 1 | 2 | 1 | 1 | 1 |
| *PTHR46315* | Spermine Synthase | 0 | 3 | 1 | 2 | 1 | 1 | 1 |
| *PTHR11360* | Monocarboxylate Transporter | 0 | 1 | 2 | 3 | 1 | 1 | 1 |
| *PTHR44442* | 3-Keto-Steroid Reductase | 0 | 3 | 1 | 1 | 1 | 1 | 1 |
| *PTHR19964* | Multiple PDZ Domain Protein | 0 | 3 | 1 | 1 | 1 | 1 | 1 |
| *PTHR15741* | Basic Helix-Loop-Helix Zip Transcription Factor | 0 | 3 | 1 | 1 | 1 | 1 | 1 |
| *PTHR42690* | Threonine Synthase Family Member | 0 | 3 | 1 | 1 | 1 | 1 | 1 |

**Table S12 Positively selected genes (PSGs) in *C. heheva***

| **No.** | **Gene ID** | **Gene Name** | **Abbreviation** | | **Omega** |
| --- | --- | --- | --- | --- | --- |
| **1** | ChiHeh_gene24148 | Exosome complex component RRP43 | EXOSC8 | 6.83705 | |
| **2** | ChiHeh_gene20072 | Isoleucine--tRNA ligase, cytoplasmic | IARS1 | 3.37011 | |
| **3** | ChiHeh_gene25641 | Phospholysine phosphohistidine inorganic pyrophosphate phosphatase | LHPP | 6.27734 | |
| **4** | ChiHeh_gene31948 | Dynactin subunit 1 | DCTN1 | 7.06391 | |
| **5** | ChiHeh_gene2856 | MutS protein homolog 4 | MSH4 | 2.92004 | |
| **6** | ChiHeh_gene15929 | Transcription factor Sox-6 | SOX6 | 9.79661 | |
| **7** | ChiHeh_gene11918 | Spinster homolog 1 | SPNS1 | 4.04511 | |
| **8** | ChiHeh_gene18484 | Microsomal triglyceride transfer protein large subunit | MTTP | 3.39416 | |
| **9** | ChiHeh_gene35223 | PAB-dependent poly(A)-specific ribonuclease subunit PAN2 | PAN2 | 5.84083 | |
| **10** | ChiHeh_gene2338 | Centromere/kinetochore protein zw10 homolog | ZW10 | 2.25404 | |
| **11** | ChiHeh_gene23426 | MAU2 sister chromatid cohesion factor | MAU2 | 6.00742 | |
| **12** | ChiHeh_gene22022 | Small RNA 2'-O-methyltransferase | HEN1 | 22.11709 | |
| **13** | CheHeh_gene5792 | Ankyrin repeat domain-containing protein 60 | ANKRD6 | 36.77157 | |
| **14** | ChiHeh_gene2437 | Uncharaterized protein |  | 5.84394 | |
| **15** | ChiHeh_gene7033 | ER degradation-enhancing alpha-mannosidase-like protein 1 | EDEM1 | 49.6507 | |
| **16** | ChiHeh_gene25677 | Serine/threonine-protein phosphatase 4 regulatory subunit 2 | PPP4R2 | 29.95628 | |
| **17** | ChiHeh_gene6288 | 18S rRNA aminocarboxypropyltransferase | TER3 | 6.98563 | |
| **18** | ChiHeh_gene21377 | U3 small nucleolar RNA-interacting protein 2 | RRP9 | 7.58009 | |
| **19** | ChiHeh_gene22931 | Diacylglycerol kinase gamma | DGKG | 1.5207 | |
| **20** | ChiHeh_gene27209 | Thioredoxin reductase 1, cytoplasmic-like | TXNRD1 | 16.1724 | |
| **21** | ChiHeh_gene15719 | Pyruvate kinase M1/2 | PKM | 3.24103 | |
| **22** | ChiHeh_gene816 | Mitochondrial-processing peptidase subunit alpha-like (LOC110987546), mRNA | PMPCA | 4.7463 | |
| **23** | ChiHeh_gene11519 | Lysine acetyltransferase 8 | KAT8 | 6.79493 | |
| **24** | ChiHeh_gene5454 | TBC1 domain family member 20-like | TBC1D20 | 62.67164 | |
| **25** | ChiHeh_gene35171 | Calpain 7 | CAPN7 | 4.89478 | |
| **26** | ChiHeh_gene9027 | Nucleolar protein 58 | NOP58 | 4.67224 | |
| **27** | ChiHeh_gene3731 | tRNA wybutosine-synthesizing protein 5-like | TYW5 | 6.51395 | |
